# Perceptual difficulty modulates the direction of information flow in familiar face recognition

**DOI:** 10.1101/2020.08.10.245241

**Authors:** Hamid Karimi-Rouzbahani, Farzad Ramezani, Alexandra Woolgar, Anina Rich, Masoud Ghodrati

## Abstract

Humans are fast and accurate when they recognize familiar faces. Previous neurophysiological studies have shown enhanced representations for the dichotomy of familiar vs. unfamiliar faces. As familiarity is a spectrum, however, any neural correlate should reflect graded representations for more vs. less familiar faces along the spectrum. By systematically varying familiarity across stimuli, we show a neural familiarity spectrum using electroencephalography. We then evaluated the spatiotemporal dynamics of familiar face recognition across the brain. Specifically, we developed a novel informational connectivity method to test whether peri-frontal brain areas contribute to familiar face recognition. Results showed that feed-forward flow dominates for the most familiar faces and top-down flow was only dominant when sensory evidence was insufficient to support face recognition. These results demonstrate that perceptual difficulty and the level of familiarity influence the neural representation of familiar faces and the degree to which peri-frontal neural networks contribute to familiar face recognition.

## Introduction

Faces are crucial for our social interactions, allowing us to extract information about identity, gender, age, familiarity, intent and emotion. Humans categorize familiar faces more quickly and accurately than unfamiliar ones, and this advantage is more pronounced under difficult viewing conditions, where categorizing unfamiliar faces often fails (Ramon and Gobbini, 2018; Young and Burton, 2018). The neural correlates of this behavioral advantage suggest an enhanced representation of familiar over unfamiliar faces in the brain (Dobs et al., 2019; Landi and Freiwald, 2017). Here, we focus on addressing two major questions about familiar face recognition. First, whether there is a “familiarity spectrum” for faces in the brain with enhanced representations for more vs. less familiar faces along the spectrum. Second, whether higher-order frontal brain areas contribute to familiar face recognition, as they do to object recognition (Bar et al,. 2006; Summerfield et al., 2006; Goddard et al., 2016; Karimi-Rouzbahani et al., 2019), and whether levels of face familiarity and perceptual difficulty (as has been suggested previously (Woolgar et al., 2011; Woolgar et al., 2015)) impact the involvement of frontal cognitive areas in familiar face recognition.

One of the main limitations of previous studies, which hinders our progress in answering our first question, is that they mostly used celebrity faces as the familiar category (Ambrus et al., 2019; Collins et al., 2018; Dobs et al., 2019). As familiar faces can range widely from celebrity faces to highly familiar ones such as family members, relatives, friends, and even one’s own face (Ramon and Gobbini, 2018), these results might not reflect the full familiarity spectrum. A better understanding of familiar face recognition requires characterizing the computational steps and representations for sub-categories of familiar faces, including personally familiar, visually familiar, famous, and experimentally learned faces. Such face categories do not only differ in terms of how much exposure the individual has had to them, but also the availability of personal knowledge, relationships, and emotions associated with the identities in question (Leppänen and Nelson, 2009; Ramon and Gobbini, 2018; Kovács, 2020). However, we still expect that potentially enhanced representations for more vs. less familiar faces, as they modulate the behavior, can also be detected using neuroimaging analysis. Moreover, these categories may vary in terms of the potential for top-down influences in the process. Importantly, while a few functional magnetic resonance imaging (fMRI) studies have investigated the differences between different levels of familiar faces (Gobbini et al., 2004; Landi and Freiwald, 2017; Leibenluft et al., 2004; Ramon et al., 2015; Sugiura et al., 2015; Taylor et al., 2009), there are no studies that systematically compare the temporal dynamics of *information processing* across this familiarity spectrum. Specifically, while event-related potential (ERP) analyses have shown amplitude modulation by levels of face familiarity (Henson et al., 2008; Kaufmann et al., 2009; Schweinberger et al., 2002; Huang et al., 2017), they remain silent about whether more familiar faces are represented more distinctly than less familiar faces - amplitude modulation does not necessarily mean that information is being represented. To address this issue, we can use multivariate pattern analysis (MVPA or decoding; Ambrus et al., 2019; Karimi-Rouzbahani et al., 2017a), which provides higher sensitivity (Norman et al., 2006) than univariate (e.g., ERP) analysis, to compare the amount of information in each of the familiarity levels.

In line with our second question, recent human studies have compared the neural dynamics for familiar versus unfamiliar face processing using the high temporal resolution of electroencephalography (EEG; Ambrus et al., 2019; Collins et al., 2018) and magnetoencephalography (MEG; Dobs et al., 2019). These studies have found that familiarity affects the initial time windows of face processing, which are generally attributed to the feed-forward mechanisms of the brain. In particular, they have explored the possibility that the face familiarity effect occurs because these faces have been seen repeatedly, leading to the development of low-level representations for familiar faces in the occipito-temporal visual system. This in turn facilitates the flow of familiar face information in a bottom-up feed-forward manner from the occipito-temporal to the frontal areas for recognition (di Oleggio Castello and Gobbini, 2015; Ramon et al., 2015; Ellis et al., 1979; Young and Burton, 2018). On the other hand, studies have also shown the role of frontal brain areas in facilitating the processing of visual inputs (Bar et al., 2006; Kveraga et al., 2007; Goddard et al., 2016; Karimi-Rouzbahani et al., 2019), such as faces (Kramer et al., 2018; Summerfield et al., 2006), by feeding back signals to the face-selective areas in the occipito-temporal visual areas, particularly when the visual input is ambiguous (Summerfield et al., 2006) or during face imagery (Mechelli et al., 2004; Johnson et al., 2007). These top-down mechanisms, which were localized in medial prefrontal cortex (MPFC), have been suggested (but not quantitatively supported) to reflect feedback of (pre-existing) face templates, against which the input faces are compared for correct recognition (Summerfield et al., 2006) in a recollection procedure (Brown and Banks, 2015). A more recent fMRI study showed that there is significant face selectivity in the inferior frontal gyrus (IFG) over the frontal cortex and that the same area is strongly connected to the well-stablished face-selective superior temporal sulcus (STS) over the temporal cortex (Davies-Thompson and Andrews, 2012), which was consistent with a previous diffusion tensor imaging study (Ethofer et al., 2011). Despite the large literature of face recognition supporting the roles of both the peri-occipital (e.g. Fusiform face area, STS) and peri-frontal^1^ (e.g. IFG, MPFC and posterior cingulate cortex (Ramon et al., 2015)) brain areas (i.e. feed-forward and feedback mechanisms), their potential interactions in familiar face recognition have remained ambiguous (see for reviews Ramon and Gobbini, 2018; Duchaine and Yovel, 2015). We develop novel connectivity methods to track the flow of information along the feed-forward and feedback mechanisms and assess the role of these mechanisms in familiar face recognition.

One critical aspect of the studies that successfully detected top-down peri-frontal to peri-occipital feedback signals (Bar et al., 2006; Summerfield et al., 2006; Goddard et al., 2016) has been the *active* involvement of the participant in a task. In recent E/MEG studies reporting support for a feed-forward explanation of the face familiarity effect, participants were asked to detect target faces (Ambrus et al., 2019) or find a match between faces in series of consecutively presented faces (Dobs et al., 2019). This makes familiarity irrelevant to the task of the participant. Such indirect tasks may reduce the involvement of top-down familiarity-related feedback mechanisms, as was demonstrated by a recent study (Kay et al., 2017), which found reduced feedback signals (from intraparietal to ventro-temporal cortex) when comparing fixation versus an active task in an fMRI study. Therefore, to answer our research questions and fully test the contribution of feedback to the familiarity effect, we need active tasks that are affected by familiarity.

Timing information is also crucial in evaluating the flows of feed-forward and feedback information as these processes often differ in the temporal dynamics. With the advent of informational connectivity analyses, we now have the potential to examine the interaction of information between feed-forward and feedback mechanisms to characterize their potential spatiotemporal contribution to familiar face recognition (Goddard et al., 2016; Goddard et al., 2019; Anzellotti and Coutanche, 2018; Basti et al., 2020; Karimi-Rouzbahani et al., 2020a). However, this requires novel methods to track the flow of familiarity information from a given brain area to a destination area and link this flow to the behavioral task goals to confirm its biological relevance. Such analyses can provide valuable insights for understanding the neural mechanisms underlying familiar face recognition in humans.

In our study, participants performed a familiar vs. unfamiliar face categorization task on sequences of images selected from four face categories (i.e., unfamiliar, famous, self, and personally familiar faces), with dynamically updating noise patterns, while their EEG data were recorded. It was crucial to use dynamic noise in this study. If stimuli were presented statically for more than ~200ms, this would result in a dominant feed-forward flow of information simply due to the incoming information (Goddard et al., 2016; Karimi-Rouzbahani, 2019; Lamme et al., 2000). On the other hand, if we present stimuli for very brief durations (e.g. < 50 ms), there may be insufficient time to evoke familiarity processing. By varying the signal-to-noise ratio of each image sequence using perceptual coherence, we were able to investigate how information for the different familiar categories gradually builds up in the electrical activity recordable by scalp electrodes, and how this relates to the amount of sensory evidence available in the stimulus (perceptual difficulty). The manipulation of sensory evidence also allowed us to investigate when, and how, feedback information flow affects familiar face recognition. Using univariate and multivariate pattern analyses, representational similarity analysis (RSA) and a novel informational connectivity analysis method, we reveal the temporal dynamics of neural representations for different levels of face familiarity.

Our results show that self and personally familiar faces lead to higher perceptual categorization accuracy and enhanced representation in the brain even when sensory information is limited while famous (visually familiar) and unfamiliar face categorization is only possible in high-coherence conditions. Importantly, our novel information flow analysis suggests that in high-coherence conditions the feed-forward sweep of face category information processing is dominant, while at lower coherence levels the exchange of face category information is consistent with feedback flow of information. The change in dominance of feedback versus feed-forward effects as a function of coherence level is consistent with a dynamic exchange of information between higher-order (frontal) cognitive and visual areas depending on the amount of sensory evidence.

## Results

We designed a paradigm to study how the stimulus- and decision-related activations for different levels of face familiarity build up during stimulus presentation and how these built-up activations relate to the amount of sensory evidence about each category. We recorded EEG data from human participants (n=18) while they categorized face images as familiar or unfamiliar. We varied the amount of sensory evidence by manipulating the phase coherence of images on different trials (Figure 1A). In each 1.2 s (max) sequence of image presentation (trial), the pattern of noise changed in each frame (16.7 ms) while the face image and the overall coherence level remained the same. Familiar face images (n=120) were selected equally from celebrity faces, photos of the participants’ own face, and personally familiar faces (e.g., friends, family members, relatives of the participant) while unfamiliar face images (n=120) were completely unknown to participants before the experiment. Within each block of trials, familiar and unfamiliar face images with different coherence levels were presented in random order.

**Figure 1.**
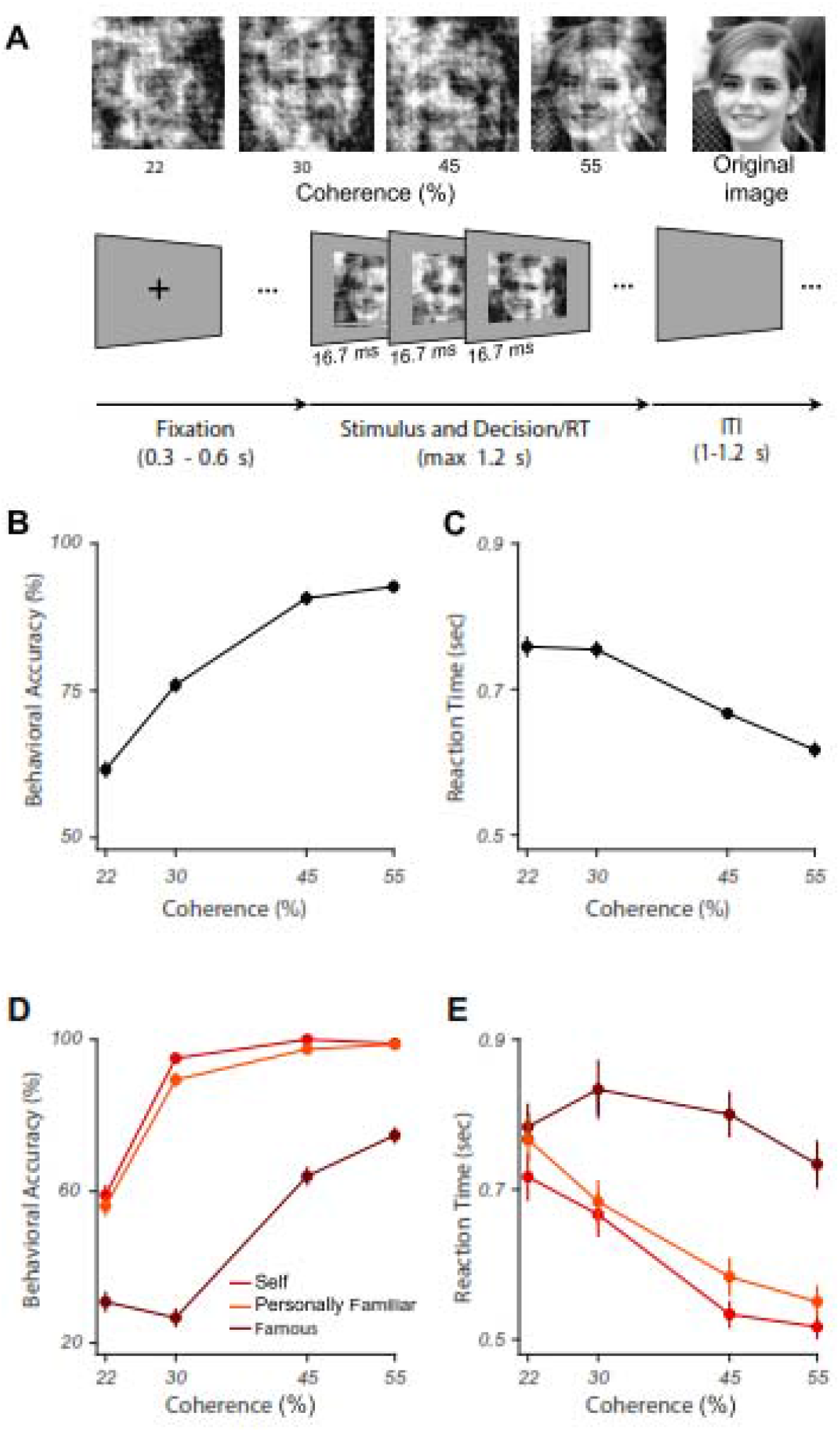
Experimental design and behavioral results for familiar vs. unfamiliar face categorization. (A) Upper row shows a sample face image (from the famous category) at the four different phase coherence levels (22, 30, 45, and 55%) used in this experiment, in addition to the original image (not used). Lower row shows schematic representation of the experimental paradigm. In each trial, a black fixation cross was presented for 300-600 ms (randomly selected). Then, a noisy and rapidly updating (every 16.7 ms) stimulus of a face image (unfamiliar, famous, personally familiar, or self), at one of the four possible phase coherence levels, was presented until response, for a maximum of 1.2 s. Participants had to categorize the stimulus as familiar or unfamiliar by pressing one of two buttons (button mappings swapped across the two sessions, counterbalanced across participants). There was then a variable inter-trial interval (ITI) lasting between 1-1.2 s (chosen from a uniform random distribution; see a demo of the task here *https://osf.io/n7b8f/*). (B) Mean behavioral accuracy for face categorization across all stimuli, as a function of coherence levels; (C) Median reaction times for correctly categorized face trials across all conditions, as a function of coherence levels. (D) and (E) show the results for different familiar face sub-categories. Error bars in all panels are the standard error of the mean across participants (smaller for panels B and C).

### Levels of face familiarity are reflected in behavioral performance

We quantified our behavioral results using accuracy and reaction times on correct trials. Specifically, accuracy was the percentage of images correctly categorized as either familiar or unfamiliar. All participants performed with high accuracy (>92%) at the highest phase coherence (55%), and their accuracy was significantly lower (~62%) at the lowest coherence (22%; F(3,272)=75.839, p<0.001; Figure 1B). The correct reaction times show that participants were significantly faster to categorize the face at high phase coherence levels than lower ones (F(3,272)=65.797, p<0.001, main effect; Figure 1C). We also calculated the accuracy and reaction times for the sub-categories of the familiar category separately (i.e. famous, personally familiar and self). The calculated accuracy here is the percentage of correct responses within each of these familiar sub-categories. The results show a gradual increase in accuracy as a function of phase coherence and familiarity (Figure 1D, two-way ANOVA. factors: coherence level and face category. Face category main effect: F(2,408)=188.708, p<0.001, coherence main effect: F(3,408)= 115.977, p<0.001, and interaction: F(6,408)=12.979, p<0.001), with the highest accuracy in categorizing their own (self), then personally familiar, and finally famous (or visually familiar) faces. The reaction time analysis also showed a similar pattern where participants were fastest to categorize self faces, then personally familiar and famous faces (Figure 1E, two-way ANOVA, factors: coherence level and face category. Face category main effect: F(2,404)=174.063, p<0.001, coherence main effect: F(3,404)= 104.861, p<0.001). We did not evaluate any potential interaction between coherence levels and familiarity levels as it does not address any hypothesis in this study. All reported p-values were corrected for multiple comparisons at p<0.05 using Bonferroni correction.

### Is there a “familiarity spectrum” for faces in the brain?

Our behavioral results showed that there is a graded increase in participants’ performance as a function of familiarity level - i.e., participants achieve higher performance if the faces are more familiar to them. In this section we address the first question of this study about whether we can find a familiarity spectrum in neural activations, using both the traditional univariate and novel multi-variate analyses of EEG.

#### Event-related potentials reflect behavioral familiarity effects

As an initial, more traditional, pass at the data, we explored how the neural responses were modulated by different levels of familiarity and coherence by averaging event-related potentials (ERP) across participants for different familiarity levels and phase coherences (Figure 2B). This is important as recent work failed to capture familiar face identity information from single electrodes (Ambrus et al., 2019). At high coherence, the averaged ERPs, obtained from a representative centroparietal electrode (CP2), where previous studies have found differential activity for different familiarity levels (Henson et al., 2008; Kaufmann et al., 2009; Huang et al., 2017), demonstrated an early, evoked response, followed by an increase in the amplitude proportional to familiarity levels. This showed that self faces elicited the highest ERP amplitude, followed by personally familiar, famous, and unfamiliar faces (Figure 2B for 55% phase coherence). This observation of differentiation between familiarity levels at later time points seems to support evidence accumulation over time, which is more pronounced at higher coherence levels where the brain had access to reliable information. This repeats previous findings showing differential activity for different levels of face familiarity after 200 ms in the post-stimulus onset window (Caharel et al., 2002; Wiese et al., 2019).

**Figure 2.**
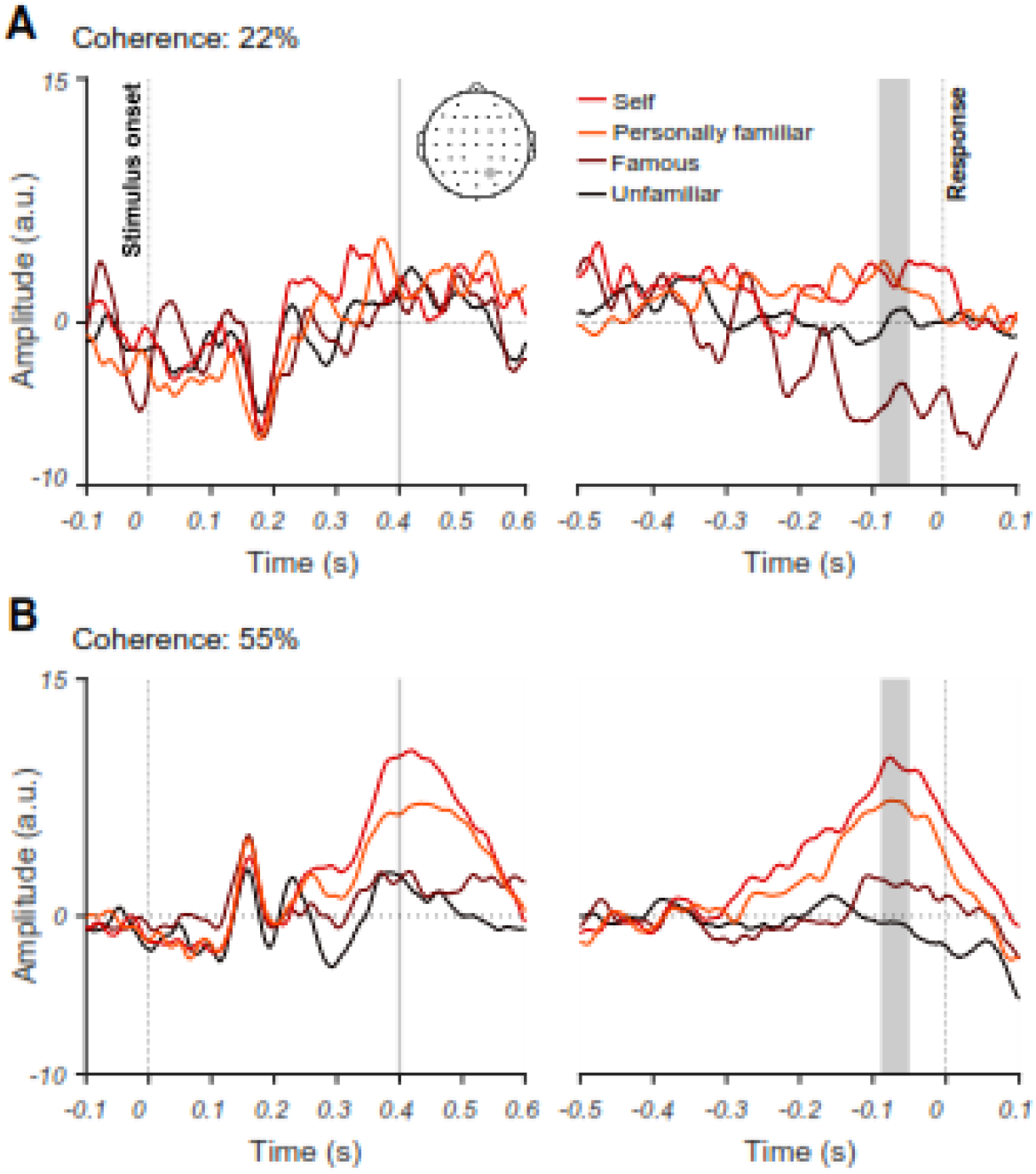
The effect of familiarity and sensory evidence on event-related potentials (ERPs). Averaged ERPs for 22% (A) and 55% (B) phase coherence levels and four face categories across all participants for an electrode at a centroparietal site (CP2). Note that the left panels show stimulus-aligned ERPs while the right panel shows response-aligned ERPs. Shaded areas show the time windows when the difference in ERPs between unfamiliar and the average of the three familiar face categories (i.e. unfamiliar-average of unfamiliar categories) were significantly (p<0.05) higher in the 55% vs. 22% coherence levels. The significance was evaluated using one-tailed independent *t*-test with correction for multiple comparisons across time at p<0.05. The differences were significant at later stages of stimulus processing around 400 ms post-stimulus onset and <100 ms before the response was given by the participant in the stimulus- and response-aligned analyses, respectively.

We also observed a similar pattern between the ERPs of different familiarity levels at the time of decision (just before the response was made). Such systematic differentiation across familiarity levels was lacking at the lowest coherence level, where the amount of sensory evidence, and behavioral performance, were low (c.f. Figure 2A for 22% phase coherence; shading areas). We observed a gradual increase in separability between the four face categories when moving from low to high coherence levels (Supplementary Figure 1). The topographic ERP maps (Supplementary Figure 2) show that the effects are not localized on the CP2 electrode, but rather distributed across the head. There are electrodes which seem to show even more familiarity information than the CP2 electrode. These results reveal the neural correlates of perceptual differences in categorizing different familiar face categories under perceptually difficult conditions.

#### Dynamics of neural representation and evidence accumulation for different face familiarity levels

Our results so far are consistent with previous event-related studies showing that the amplitude of ERPs is modulated by the familiarity of the face (Henson et al., 2008; Kaufmann et al., 2009; Schweinberger et al., 2002; Huang et al., 2017). However, more modulation of ERP amplitude does not necessarily mean enhanced representation. Moreover, we observed that the familiarity effects were distributed across the head rather than localized only on the individual CP2 electrode (Supplementary Figure 2). Therefore, looking at individual electrodes might overlook the true temporal dynamics of familiarity information, which may involve widespread brain networks (Ramon and Gobbini, 2018; Duchaine and Yovel, 2015). Here we used multivariate pattern and representational similarity analyses on these EEG data to quantify the time course of familiar vs. unfamiliar face processing. Compared to traditional single-channel (univariate) ERP analysis, MVPA allows us to capture the whole-brain widespread and potentially subtle differences between the activation dynamics of different familiarity levels (Ambrus et al., 2019; Dobs et al., 2019). Specifically, we asked: (1) how the representational dynamics of stimulus- and response-related activations change depending on the level of face familiarity; and (2) how manipulation of sensory evidence (phase coherence) affects neural representation and coding of different familiarity levels.

To obtain the temporal evolution of familiarity information across time, at each time point we trained the classifier to discriminate between familiar and unfamiliar faces. Note that the mapping between response and fingers were swapped from the first session to the next (counterbalanced across participants) and the data were collapsed across the two sessions for all analyses, which ensures the motor response cannot drive the classifier. We trained the classifier using 90% of the trials and tested it on the left-out 10% of trials using a standard 10-fold cross-validation procedure (see *Methods*). This analysis used only correct trials. Our decoding analysis shows that, up until ~200 ms after stimulus onset, decoding accuracy is near chance for all coherence levels (Figure 3A). The decoding accuracy then gradually increases over time and peaks around 500 ms post-stimulus for the highest coherence level (55%) but remains around chance for the lower coherence level (22%, Figure 3A). The accuracy for intermediate coherence levels (30% and 45%) falls between these two bounds but only reaches significance above chance for the 45% coherence level. This ramping up temporal profile suggests an accumulation of sensory evidence in the brain across the time course of stimulus presentation, which has a processing time that depends on the strength of the sensory evidence (Hanks and Summerfield, 2017; Philiastides et al., 2006).

**Figure 3.**
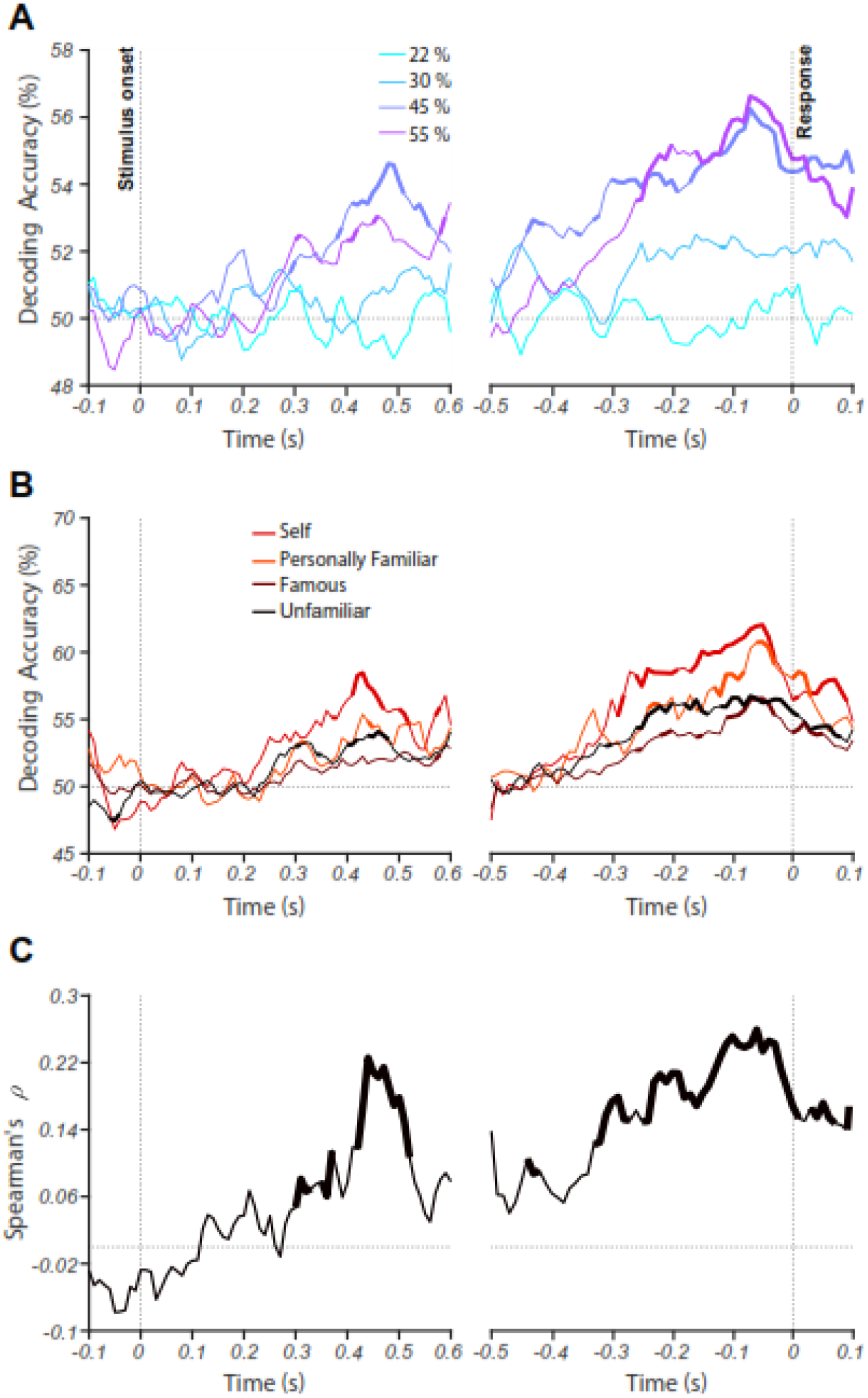
Decoding of face familiarity from EEG signals. (A) Time course of decoding accuracy for familiar versus unfamiliar faces from EEG signals for four different phase coherence levels (22%, 30%, 45%, and 55%). (B) Time course of decoding accuracy for four face categories (i.e., unfamiliar, famous, self and personally familiar faces) from EEG signals at the 55% coherence level. The chance accuracy is 50%. Thickened lines indicate the time points when the accuracy was significantly above chance level (sign rank test, FDR corrected across time, p<0.05). (C) Correlation between behavioral performance and decoding accuracy (across all conditions) over time. Thickened lines indicate the time points when the correlation was significant. The left panels show the results for stimulus-aligned analysis while the right panels show the results for response-aligned analysis (averaged over 18 participants).

After verifying that we could decode the main effect of familiarity, we turned to the first main question of the study. To examine if neural decoding could reveal the spectrum of familiarity which we observed in behavior and ERPs, we separately calculated the decoding accuracy for each of the sub-categories of familiar faces (Figure 3B): unfamiliar, famous, self and personally familiar faces (on the 55% coherence level, which showed the highest decoding in Figure 3A). The decoding accuracy was highest for self faces, both for stimulus- and response-aligned analyses, followed by personally familiar, famous and unfamiliar faces. Accuracy for the response-aligned analysis shows that the decoding gradually increased to peak decoding ~100 ms before the response was given by participants. This temporal evolution of decoding accuracy begins after early visual perception and rises in proportion to the amount of the face familiarity.

To rule out the possibility that an unbalanced number of trials in the sub-categories of familiar faces could lead to the difference in decoding accuracies between the sub-categories, we also repeated the decoding analysis by classifying each familiar subcategory from the unfamiliar category (after equalizing the number of trials across the familiar and unfamiliar categories and also across the three familiar categories), which provided similar results. We also repeated the same analysis for lower coherence levels: only the two high-coherence conditions (i.e. 45% and 55%), but not the low-coherence conditions (i.e. 22% and 30%), showed significantly above-chance decoding for all familiarity conditions (Supplementary Figure 3).

Low-level stimulus differences between conditions could potentially drive the differences between categories observed in both ERP and decoding analyses (e.g., familiar faces being more frontal than unfamiliar faces, leading to images with brighter centers and, therefore, separability of familiar from unfamiliar faces using central luminance of images; Dobs et al., 2019; Ambrus et al., 2019). To address such potential differences, we carried out a supplementary analysis using RSA (Supplementary Text and Supplementary Figures 4 and 5), which showed that such differences between images do not play a major role in the differentiation between categories.

To determine whether the dynamics of decoding during stimulus presentation are associated with tfhe perceptual task, as captured by our participants’ behavioral performance, we calculated the correlation between decoding accuracy and perceptual performance. For this, we calculated the correlation between 16 data points from decoding accuracy (4 face categories × 4 phase coherence levels) and their corresponding behavioral accuracy rates, averaged over participants. The correlation peaked ~500 ms post-stimulus (Figure 3C), which was just before the response was given. This is consistent with an evidence accumulation mechanism determining whether to press the button for ‘familiar’ or ‘unfamiliar’, which took another ~100 ms to turn into action (finger movement).

### Do higher-order peri-frontal brain areas contribute to familiar face recognition?

In this section we address the second question of this study about whether peri-frontal brain areas contribute to the recognition of familiar faces in the human brain using a novel informational connectivity analyses on EEG.

#### Task difficulty and familiarity level affect information flow across the brain

We investigated how the dynamics of feed-forward and feedback information flow changes during the accumulation of sensory evidence and the evolution over a trial of neural representations of face images. We developed a novel connectivity method based on RSA to quantify the relationships between the evolution of information based on peri-occipital EEG electrodes and those of the peri-frontal electrodes. As an advantage to previous Granger causality methods (Goddard et al., 2016; Goddard et al., 2019; Karimi-Rouzbahani et al., 2019; Kietzman et al., 2019), the connectivity method developed here allowed us to check whether the transferred signals contained *specific aspects of stimulus information*. Alternatively, it could be the case that the transferred signals might carry highly abstract but irrelevant information between the source and destination areas, which can be incorrectly interpreted as connectivity (Anzellotti and Coutanche, 2018; Basti et al., 2020). Briefly, feed-forward information flow is quantified as the degree to which the information from peri-occipital electrodes at present time contributes to the information recorded at peri-frontal electrodes at a later time point, which reflects moving the frontal representation closer to that required for task goals. Feedback flow is defined as the opposite: the contribution to information at peri-frontal electrodes at the present time to that recorded later at peri-occipital electrodes at a later time point (Figure 4A).

**Figure 4.**
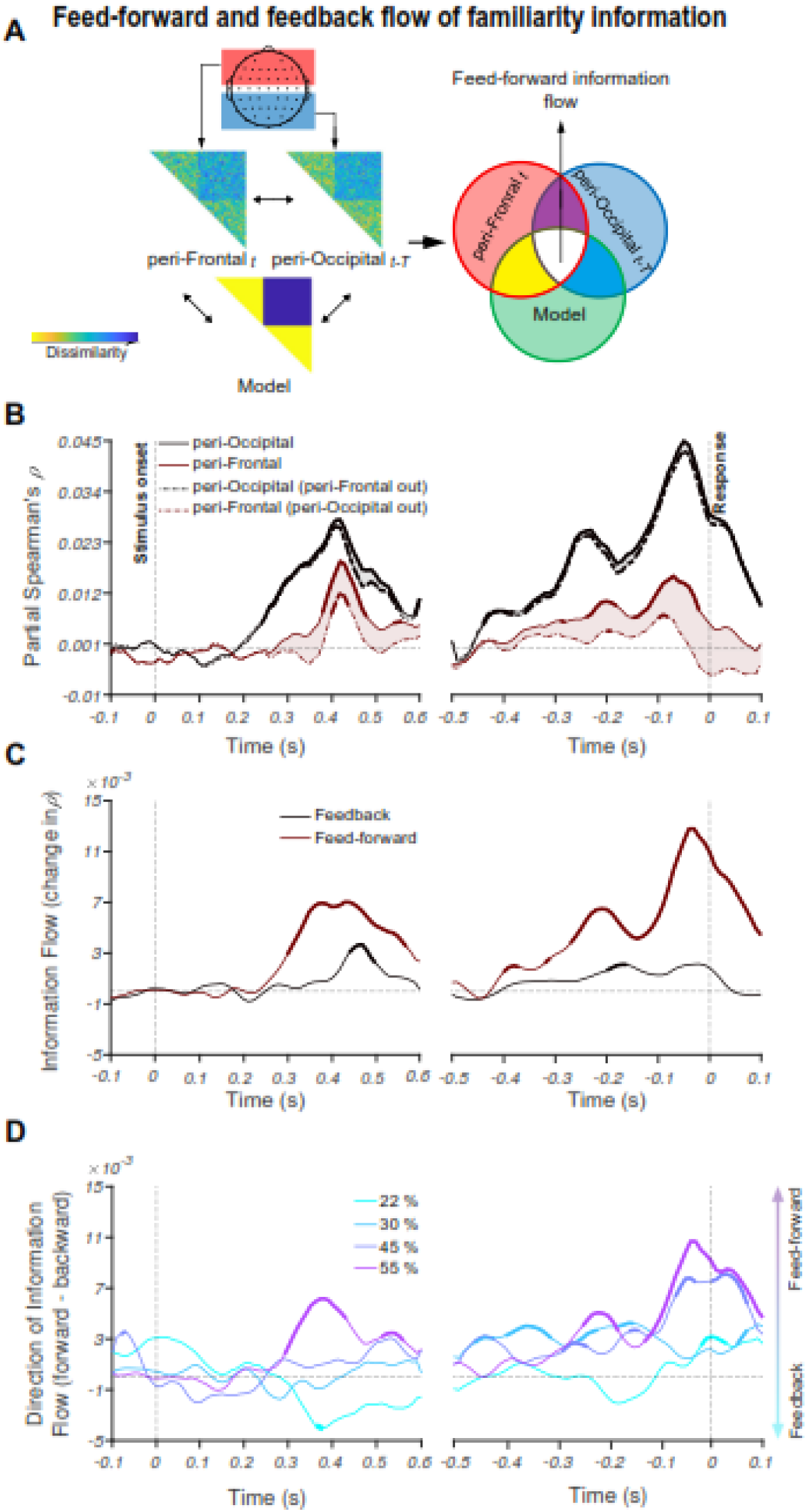
Feed-forward and feedback information flow revealed by RSA. **(A)** A schematic presentation of the method for calculating informational connectivity between the peri-frontal and peri-occipital electrodes, termed feed-forward and feedback information flow. Feed-forward information flow is calculated as difference of the correlation between the present time peri-frontal neural RDM and the predicted model RDM and the same correlation when the earlier peri-occipital neural RDM is partialled out from it. This is shown in the Venn diagram on the right. The summation of white and yellow areas reflect the correlation between the peri-frontal and the model RDMs while the yellow area reflects the same correlation after partialling out the peri-occipital area at the earlier time point. The difference between the two (i.e. white = (white + yellow)– yellow) is considered to be feed-forward flow of information captured by the model. Delay time (T) is 30ms. **(B)** Time course of partial Spearman’s correlations representing the partial correlations between the peri-occipital (black) and peri-frontal (brown) EEG electrodes and the model (familiar-unfamiliar model, see the inset in A) while including (solid) and excluding (dashed) the effect of the other area at phase coherence of 55%. The shaded area shows the decline in partial correlation of the current area with the model after excluding (partialling out) the RDM of the other area. Note that in both the dashed and solid lines, the low-level image statistics are partialled out of the correlations, so we call them partial correlations in both cases. **(C)** Feedforward (brown) and feedback (black) information flows obtained by calculating the value of the shaded areas in the corresponding curves in B. **(D)** Direction of information flow for different coherence levels, determined as the difference between feed-forward and feedback information flow showed in C. Thickened lines indicate time points at which the difference is significantly different from zero (sign permutation test and corrected significance level at p□<□0.05), and black dotted lines indicate 0 correlation. The left panels show the results for stimulus-aligned analysis while the right panels represent the results for response-aligned analysis.

The results show that at the highest coherence level (55%), information flow is dominantly in the feed-forward direction. This is illustrated by the shaded area in Figure 4B where partialling out the peri-frontal from peri-occipital correlations only marginally reduces the total peri-occipital correlation (Figure 4B, black curves and shaded area), meaning that there is limited information transfer from peri-frontal to peri-occipital. In contrast, partialling out the peri-occipital from peri-frontal correlations leads to a significant reduction in peri-frontal correlation, reflecting a feed-forward transfer of information (Figure 4B, brown curves and shaded area). This trend is also seen for response-aligned analysis.

These differences are shown more clearly in Figure 4C where the peaks of feed-forward and feedback curves show that the feed-forward information is dominant earlier, followed by feedback information flow, as shown by the later peak of feedback dynamics. These results suggest that when the sensory evidence is high, feed-forward information flow may be sufficient for categorical representation and decision making while feedback only slightly enhances the representation. However, in lower coherence levels (i.e., low sensory evidence), the strength of information flow is either equivalent between feed-forward and feedback directions (30%, 45%) or dominantly feedback (22%, Figure 4D).

Here, we can see that the lower sensory evidence correlates with greater engagement of feedback mechanisms, suggesting that feedback is recruited to boost task-relevant information in sensory areas under circumstances where the input is weak. This is consistent with the dynamics and relative contribution of feedback and feed-forward mechanisms in the brain varying with the sensory evidence / perceptual difficulty of the task.

Importantly, we also were interested in whether the degree of familiarity changes the direction of information flow between the peri-frontal and peri-occipital brain areas. For this analysis, we collapsed the data across all coherence levels to look specifically at the impact of face familiarity. We generated specific RDM models to evaluate how much information about unfamiliar faces vs. all unfamiliar faces as a group (Figure 5A) and each subcategory of familiar faces (i.e., famous, personally familiar and self; Figure 5B) individually were transferred between the two brain areas. To avoid any bias from a different number of elements in the RDM matrices, we only compared equal-sized conditions and present the results in separate panels (i.e. familiar vs. unfamiliar (Figure 5A) and subcategories of familiar faces (Figure 5B)). While the unfamiliar category showed a non-significant flow in either direction, the familiar category showed significant feed-forward flow of information in the stimulus-aligned data starting from 300 ms post-stimulus onset (Figure 5A). Among the familiar sub-categories, only the self category showed significant feed-forward information flow starting to accumulate after the stimulus onset, reaching sustained significance ~500 ms. The less familiar categories of famous and personally familiar did not reach significance. In the response-aligned analysis, again, the significant time points show the domination of feed-forward flow for the familiar category (Figure A) but not the unfamiliar category, and the self category but not the other sub-categories of familiar faces (Figure 5B). Together, these results suggest that while the information about the unfamiliar category did not evoke a particular dominance of information flow in either direction, the representations of familiar and self faces showed dominant feed-forward information flow from the peri-occipital to the peri-frontal brain areas. Note that, in this analysis, we also tried to minimize the effect of the participants’ decision and motor response in the models by excluding the opposing category (i.e. unfamiliar category when evaluating the familiar models and *vice versa*), which could have potentially contributed to the information flows in the previous analysis (c.f. Figure 4).

**Figure 5.**
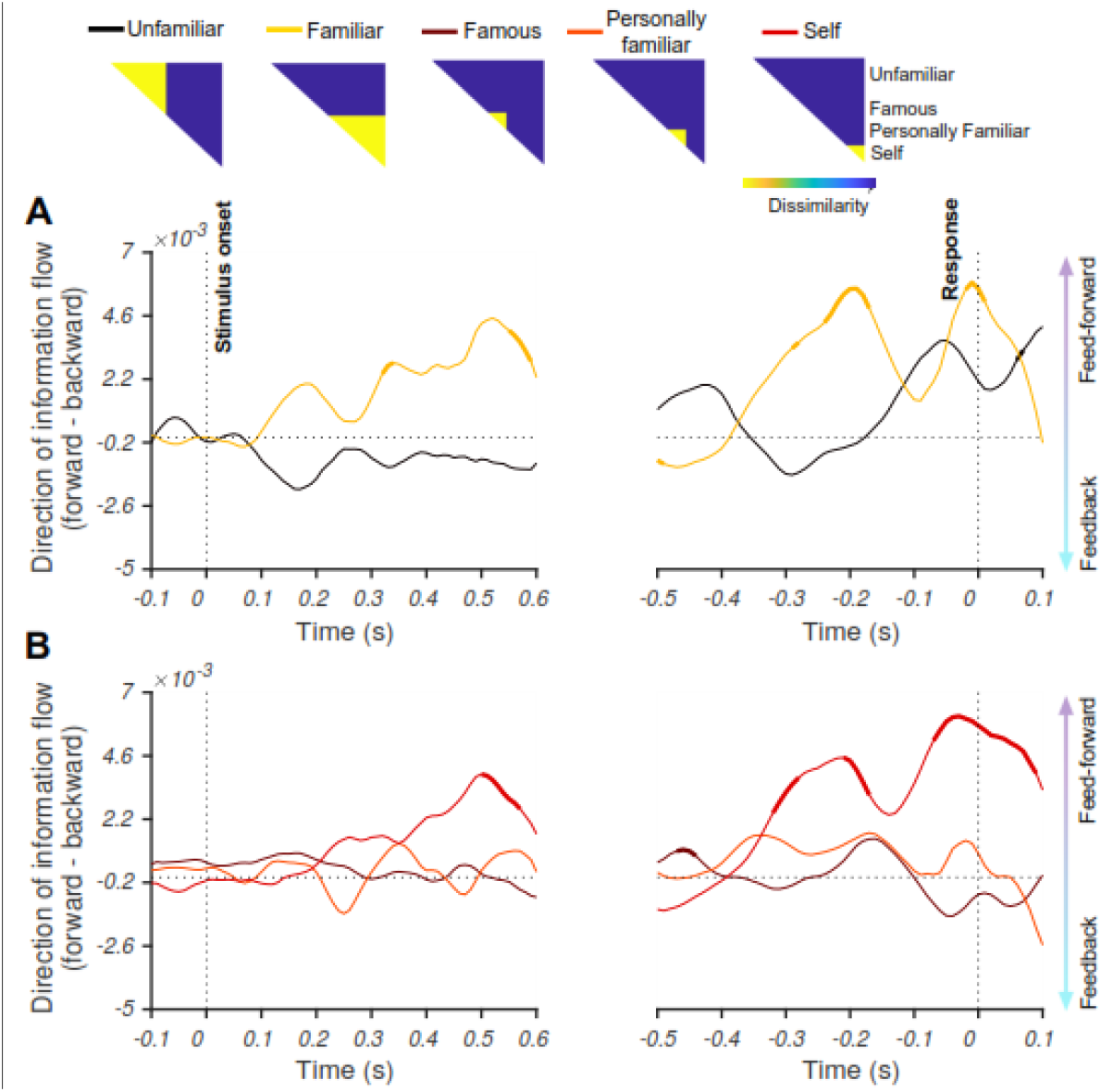
Directions of information flow for familiar, unfamiliar and different levels of familiarity. The models, as depicted on the top, are constructed to measure the extent and timing by which information about unfamiliar and familiar (A), and each familiar sub-category (B) moves between the peri-occipital and peri-frontal brain areas. Feed-forward information flow is calculated as difference of the correlation between the present time peri-frontal neural RDM and the predicted model RDM and the same correlation when the earlier peri-occipital neural RDM is partialled out from it. This is shown in the Venn diagram in Figure 4A. The summation of white and yellow areas reflect the correlation between the peri-frontal and the model RDMs while the yellow area reflects the same correlation after partialling out the peri-occipital area at the earlier time point. The difference between the two (i.e. white = (white + yellow)– yellow) is considered to be feed-forward flow of information captured by the model. Delay time (T) is 30ms. The yellow areas in the models refer to the target category (including unfamiliar, familiar, famous, personally familiar and self faces). Thickened lines indicate time points at which the difference is significantly different from zero (sign permutation test and corrected for multiple comparisons at significance level of p□<□0.05), and black horizontal dotted lines indicate 0 correlation. The left panel shows the result for stimulus-aligned analysis while the right panels represent the result for response-aligned analysis.

Together, the results of the information connectivity analysis suggest that, in familiar face recognition, both top-down and bottom-up mechanisms play a role, with the amount of sensory evidence determining their relative contribution. It also suggests that the degree to which sensory information is processed feed-forward can be modulated by the familiarity level of the stimulus.

## Discussion

This study investigated the neural mechanisms of familiar face recognition. We asked how familiarity affected the contribution of feed-forward and feedback processes in face processing. We first showed that manipulating the familiarity affected the informational content of neural responses about face category, in line with a large body of behavioral literature showing an advantage of familiar over unfamiliar face processing in the brain. Then, we developed a novel method of informational connectivity analysis to track the exchange of familiarity information between peri-occipital and peri-frontal brain areas to see if frontal brain areas contribute to familiar face recognition. Our results showed that when the perceptual difficulty was low (high sensory evidence), the flow of face familiarity information was consistent with a feed-forward account. On the other hand, when the perceptual difficulty was high (low sensory evidence), the dominant flow of face familiarity information reversed, which we interpret as reliance on feedback mechanisms. Moreover, when teasing apart the effect of task and response from neural representations, only the familiar faces, but not the unfamiliar faces, showed a dominance of feed-forward flow of information, with maximum flow for the most familiar category, the self faces.

Our results are consistent with the literature suggesting that visual perception comprises both feed-forward and feedback neural mechanisms transferring information between the peri-occipital visual areas and the peri-frontal higher-order cognitive areas (Bar et al., 2006; Summerfield et al., 2006; Goddard et al., 2016; Karimi-Rouzbahani et al., 2017b; Karimi-Rouzbahani et al., 2017c; Karimi-Rouzbahani et al., 2019). However, previous experimental paradigms and analyses did not dissociate feedback and feed-forward information flow in familiar face recognition, and argued for a dominance of feed-forward processing (Dobs et al., 2019; di Oleggio Castello and Gobbini, 2015; Ellis et al., 1979; Young and Burton, 2018). The more nuanced view we present is important because stimulus familiarity, similar to other factors including levels of categorization (superordinate vs. basic level; Besson et al., 2017; Praß et al., 2013), task difficulty (Chen et al., 2008; Woolgar et al., 2015; Kay et al., 2017) and perceptual difficulty (Fan et al., 2020; Hupe et al., 1998; Gilbert and Li, 2013; Gilbert and Sigman, 2007; Lamme and Roelfsema, 2000; Woolgar et al., 2011), may affect the complex interplay of feed-forward and feedback mechanisms in the brain.

Our results showed that the contribution of peri-frontal to peri-occipital feedback information was inversely proportional to the amount of sensory evidence about the stimulus. Specifically, we only observed feedback when the sensory evidence was lowest (high perceptual difficulty) in our face familiarity categorization task. Although a large literature has provided evidence for the role of top-down feedback in visual perception, especially when sensory visual information is low, they generally evaluated the feedback mechanisms within the visual system (Ress et al., 2000; Lamme and Roelfsema, 2000; Super et al., 2001; Lamme et al., 2002; Pratte et al., 2013; Fenske et al., 2006; Lee and Mumford, 2003; Felleman et al., 1991; Delorme et al., 2004; Mohsenzadeh et al., 2018; Kietzmann et al., 2019) rather than across the fronto-occpital brain networks (Bar et al., 2006; Summerfield et al., 2006; Goddard et al., 2016; Karimi-Rouzbahani et al., 2018; Karimi-Rouzbahani et al., 2019). Our findings support theories suggesting that fronto-occipital information transfer may feedback (pre-existing) face templates, against which the input faces are compared for correct recognition (Bar et al., 2006; Summerfield et al., 2006). Previous results could not determine the content of the transferred signals (Bar et al., 2006; Summerfield et al., 2006; Goddard et al., 2016; Karimi-Rouzbahani et al., 2018; Karimi-Rouzbahani et al., 2019). Here, using our novel connectivity analyses, we showed that the transferred signal contained information which contributed to the categorization of familiar and unfamiliar faces.

Despite methodological differences, our findings support previous human studies showing increased activity in lower visual areas when the cognitive and perceptual tasks were difficult relative to easy, which the authors attributed to top-down contributions (Ress et al., 2000; Kay et al., 2017). However, due to the low temporal resolution of fMRI, these studies cannot show the temporal evolution of these top-down contributions or the validity of the hypothesized direction. Importantly, the observed increase in activity in lower visual areas does not necessarily correspond to the enhancement of neural representations in those areas - increased univariate signal does not show whether there is more information that will support performance. Electrophysiological studies in animals have also shown that cortical feedback projections robustly modulate responses of early visual areas when sensory evidence is low, or the stimulus is difficult to segregate from the background figure (Hupe et al., 1998). A recent study has also found cortical feedback modulated the activity of neurons in the dorsolateral geniculate nucleus (dLGN), which was less consistent when presenting simple vs. complex grating stimuli (Spacek et al., 2019). Therefore, varying perceptual difficulty seems to engage different networks and processing mechanisms, and we show here that this also pertains to faces: less difficult stimuli such as our high-coherence faces seem to be predominantly processed by the feed-forward mechanisms, while more difficult stimuli such as our low-coherence faces recruit both feed-forward and feedback mechanisms. However, the exact location of the feedback in all these studies, including ours, remains to be determined with the development of more accurate modalities for neural activity recording.

We observed that the direction of information flow is influenced by the familiarity of the stimulus. The models of familiar faces and self faces, evoked a dominant flow of feed-forward information. The unfamiliar category, however, did not evoke information flow in any direction, as evaluated by our connectivity method. This is consistent with enhanced representations of familiar face categories in the feed-forward pathways (Dobs et al., 2019; di Oleggio Castello and Gobbini, 2015; Ellis et al., 1979; Young and Burton, 2018), which, in turn, requires less top-down contributions to facilitate the perception of relevant information (Bar et al., 2006; Gilbert and Sigman, 2007). Our results might initially seem inconsistent with Fan et al.’s (2020) study, which did not report significant differences between the temporal dynamics of familiar and unfamiliar face representations; however, they only used famous faces within the familiar face spectrum. In our sub-category analysis, we also did not observe differences between famous faces and unfamiliar faces; our main findings were from highly familiar self faces. Overall, then, our results suggest that processing of familiar faces, especially the most familiar (self) faces, is dominated by feed-forward information flow.

One assumption in the connectivity analysis of the current work, as in many previous ones (Goddard et al., 2016; Clarke et al., 2018; Kietzman et al., 2019), is that all categories of faces used here involve neural mechanisms from both the peri-frontal and peri-occipital areas. However, this is not necessarily the case; we know from the face recognition literature that peri-frontal brain areas (as defined in this study) play role in the processing of face-relevant information such as social, dynamic and eye-movement-related aspects in cooperation with superior temporal brain areas (Duchaine and Yovel, 2015; superior temporal areas are grouped here in the peri-occipital category). On the other hand, the peri-occipital brain areas have been suggested to dominantly process lower order sensory-level face features with relatively more independence from peri-frontal brain areas (Collins and Olsen, 2014). This suggests that our connectivity analysis might provide a stronger flow for one aspect of information than the other depending on the potentially distinct neural network involved for each. However, to the best of our knowledge, no studies have suggested distinct networks for the processing of the conditions which we compared (familiar vs. unfamiliar faces or familiarity levels). Thus, we cannot rule out the possibility that there might be factors attributable to a subset of categories, but not others, that involve distinct networks. For example, it could be the case that familiar faces, but not unfamiliar ones, involve emotion networks which span from the posterior to the anterior brain areas (Leppänen and Nelson, 2009). To avoid this potential influence, we selected images for both familiar and unfamiliar categories with variable emotional content, but any emotional content associated with basic familiarity could not be avoided. At this point, we interpret our results as an interaction between feed-forward and feedback sweeps of information within a network, but acknowledge the potential contribution of additional frontal areas for one category over another.

Our results suggest that processing differs considerably for highly familiar faces. This may be because expectation and prediction play a role in (Ramon and Gobbini, 2018; Summerfield and Egner, 2009), and can potentially affect the contribution of feedback neural mechanisms in face detection (Summerfield et al., 2006). Specifically, familiar faces are generally more limited in number compared to unfamiliar faces, which can potentially make the former more predictable. However, according to the earlier visual recognition literature (Bar et al., 2006; Summerfield et al., 2006), if anything, this would have evoked more pronounced feedback signals for the familiar faces vs. unfamiliar faces in this study. In contrast to this prediction, our results showed dominant feed-forward flow of information for familiar faces, and no significant flow in either direction for unfamiliar faces. Therefore, it seems unlikely that the potential difference in expectation between familiar and unfamiliar categories could explain our information flow results.

Results also show that, in lower coherence levels, the information about the familiarity levels was generally stronger than the information about familiarity itself (as captured by familiar-unfamiliar model RDM; Supplementary Figure 4). This suggests a lower threshold for the appearance of familiarity level compared to familiar-unfamiliar representations, which are differentially developed through life-time experience and task instructions, respectively. Specifically, development of neural representations reflecting familiarity levels could be a result of exposure to repetitive faces, which can lead to developing face-specific representations in the visual system (Dobs et al., 2019), while task instructions could temporarily enhance the processing of relevant information in the brain through top-down mechanisms (Hebart et al., 2018; Karimi-Rouzbahani et al., 2019). This is probably the reason for the dominance of feedback information flow in the processing of familiarity information (Figure 5A).

The RSA-based connectivity method used in this study follows a recent shift towards informational brain connectivity methods (Anzellotti and Coutanche, 2018; Basti et al., 2020; Keitzmann et al., 2019; Goddard et al., 2016; Clarke et al., 2018; Karimi-Rouzbahani, 2018; Karimi-Rouzbahani et al., 2019; Karimi-Rouzbahani et al., 2020a), and introduces a few distinct features compared to previous methods of connectivity analyses. Specifically, traditional connectivity methods examine inter-area interactions through indirect measures such as gamma-band synchronization (Gregoriou et al., 2009), shifting power (Bar et al., 2006) or causality in the activity patterns (Summerfield et al., 2006; Fan et al., 2020). Such connectivity methods consider simultaneous (or time-shifted) correlated activations of different brain areas as connectivity, but they are unable to examine how (if at all) relevant information is transferred across those areas. Goddard et al. (2016) developed an RSA-based Granger connectivity method to solve this issue, which allowed us and others to track the millisecond transfer of stimulus information across peri-frontal and peri-occipital brain areas (Karimi-Rouzbahani, 2018; Karimi-Rouzbahani et al., 2019; Goddard et al., 2019). This was followed by another informational connectivity method, which was similar but used regression instead of correlation in implementation (Keitzmann et al., 2019). While informative, these methods, are silent about what aspects of the representation are transferred and modulated. In other words, we need new methods to tell how (if at all) the transferred information is contributing to the representations in the destination area. Not having access to the transferred contents could lead to incorrect interpretations of connectivity for one main reason: we would not be able to tease apart transactions of distinct types of information across areas (e.g. familiar-unfamiliar discrimination, or different levels of familiarity). To address this issue, one could simply calculate the correlation between the neural and model RDMs from the source and destination areas at every time point and then calculate the Granger causality between the two time-courses of correlations. This is exactly how Clarke et al., (2018) incorporated RDM models into their connectivity to track specific aspects of the transferred information. However, this last method loses the temporal dynamics of information flow in the calculation of Granger causality, and only provides the direction of information flow. Our method circumvents this limitation (i.e. lack of temporal information) by making use of the high-dimensional representational space of the RDMs in the source and destination areas for the calculation of inter-area and area-model relationship leaving the time samples available for the evaluation of the temporal dynamics. Our method allows us to explicitly determine the content (using model RDMs), the direction (using delayed time samples across areas) and the temporal evolution (using temporally-resolved analysis) of the information transferred from the peri-frontal to peri-occipital areas and vice versa. The relevance of the transferred information is determined by the amount that the representations in the destination area are shifted towards our predefined predicted RDM models. In this way, we could determine the temporal dynamics of the contributory element of the transferred information. Despite the specificity that the model-based methods (including our proposed one) provide about the content of the transferred information, such model-based methods have the characteristic to ignore other model-irrelevant aspects of information which might be similarly or distinctly represented in the source and destination areas. In other words, while the source and destination areas might show high levels of connectivity through the “lens” of the model used, their representational geometry (as evaluated here using RDMs) might be very distinct from one another when directly compared or vice versa. Therefore, the results of model-based connectivity methods do not make any prediction about the direction and the amount of potential connectivity when using model-free connectivity methods.

Despite the advantage that informational connectivity methods provide over conventional univariate connectivity methods, further investigations (using simulated datasets with known ground-truth of information flow) are needed to fully uncover their characteristics and potential limitations. As an initial step in that direction, we simulated a simplified well-controlled dataset and applied our connectivity analysis to it to check if it could detect the imposed information flow between our simulated source and destination areas (Supplementary Figure 6). Results showed that our connectivity analysis detected correct direction and temporal dynamics of the simulated information flow. Despite this successful simulation, a full mathematical and analytical investigation will need to be performed to compare the available and the proposed informational connectivity analyses in the future.

Our results specify the neural correlates for the behavioral advantage in recognizing more vs. less familiar faces in a “familiarity spectrum”. As in previous studies, our participants were better able to categorize highly familiar than famous or unfamiliar faces, especially in low-coherence conditions (Kramer et al., 2018; Young and Burton, 2018). This behavioral advantage could result from long-term exposure to variations of personally familiar faces under different lighting conditions and perspectives, which is usually not the case for famous faces.

Our neural decoding results quantified a neural representational advantage for more familiar faces compared to less familiar ones (i.e. higher decoding for the former than the latter) to suggest that more familiar faces also lead to more distinguishable neural representations. Decoding accuracy was also proportional to the amount of sensory evidence: the higher the coherence levels, the higher the decoding accuracy. We observed that the decoding accuracy “ramped-up” and reached its maximum ~100 ms before participants expressed their decisions using a key press. These results are suggestive of sensory evidence accumulation and decision making processes during face processing in humans, consistent with previously reported data in monkey and recent single-trial ERP studies (Kelly et al., 2013; Hanks and Summerfield, 2017; Philiastides et al., 2006; Philiastides and Sajda, 2006; Shadlen and Newsome, 2001).

There was a significant correlation between MVPA accuracy and our behavioral results, showing a relationship between neural representation and behavioral outcomes. While it would be ideal to see perfect correlation between neural data and behavior, it is not usually the case (Dobs et al., 2019), which may reflect several reasons including the noise in the neural data and sub-optimal decoding of the neural codes (Karimi-Rouzbahani et al., 2020b) and/or possible non-linear relationships between neural data and behavior. In our study, while there was a difference between the neural data from personally familiar and self faces (c.f. Figures 2 and 3), there was no detectable difference in behavior, potentially reflecting a ceiling effect for both categories in above-chance conditions (i.e. coherence levels > 30%). Despite this, the correlation was significant across the four familiarity × four coherence level conditions overall during time windows later in the trial and immediately before the response. This suggests that the behavioral advantages of self and familiar faces and/or having higher sensory evidence (highest coherence) could have been driven by the enhanced neural representations.

Previous studies generally show face familiarity modulation during early ERP components such as N170, N250, and P300 (Dobs et al., 2019; Ambrus, 2019; Fan et al., 2020; Henson et al., 2008; Kaufmann et al., 2009; Schweinberger et al., 2002; Huang et al., 2017). In contrast, the time course of our EEG results showed their maximum effects after 400 ms post-stimulus onset. However, these studies typically use event-related paradigms, which evoke initial brain activations peaking at around 200 ms, whereas our dynamic masking paradigm releases the information gradually along the time course of the trial. Moreover, the extended (>200ms) static stimulation used in previous studies has been suggested to bias towards domination of feed-forward processing (Goddard et al., 2016; Karimi-Rouzbahani, 2018), because of the co-processing of the incoming sensory information and the recurrence of earlier windows of the same input (Kietzmann et al., 2019; Mohsenzadeh et al., 2018), making it hard to measure feedback. Our paradigm, while providing a delayed processing profile compared to previous studies, avoids this and also slows down the process of evidence accumulation so that it becomes more trackable in time. This does mean, however, that our time courses are not really comparable with previous ERP results.

In conclusion, our study demonstrates that the processing of face information involves both feed-forward and feedback flow of information in the brain, and which predominates depends on the strength of incoming perceptual evidence and the familiarity of the face stimulus. Our novel extension of multivariate connectivity analysis methods allowed us to disentangle feed-forward and feedback contributions to familiarity representation. This connectivity method can be applied to study a wide range of cognitive processes, wherever information is represented in the brain and transferred across areas. We also showed that the behavioral advantage for familiar face processing is robustly reflected in neural representations of familiar faces in the brain and can be quantified using multivariate pattern analyses. These new findings and methods emphasize the importance of, and open new avenues for, exploring the impact of different behavioral tasks on the dynamic exchange of information in the brain.

## Materials and Methods

### Participants

We recorded from 18 participants (15 male, aged between 20-26 years, all with normal or corrected-to-normal vision). Participants were students from the Faculty of Mathematics and Computer Science at the University of Tehran, Iran. All participants voluntarily participated in the experiments and gave their written consent prior to participation. All experimental protocols were approved by the ethical committee of the University of Tehran. All experiments were carried out in accordance with the guidelines of the Declaration of Helsinki.

### Stimuli

We presented face images of four categories, including unfamiliar, famous, self and personally familiar faces. The unfamiliar faces (n=120) were unknown to participants. The famous faces (n=40) were pictures of celebrities, politicians, and other well-known people. These faces were selected from different, publicly available face databases^2^. In both categories, half of the images were female, and half were male. To ensure that all participants knew the famous face identities, participants completed a screening task prior to the study. In this screening, we presented them with the names of famous people in our data set and asked if they were familiar with the person.

The personally familiar faces were selected from participants’ family, close relatives, and friends (n=40); self-images were photographs of participants (n=40). The images of self and personally familiar faces were selected to have varied backgrounds and appearances. On average, we collected n=45 for personally familiar and n=45 for self faces for every individual participant. All images were cropped to have 400×400 pixels and were converted to greyscale (Figure 1A). We ensured that spatial frequency, luminance, and contrast were equalized across all images. The magnitude spectrum of each image was adjusted to the average magnitude spectrum of all images in our database^3^.

The phase spectrum was manipulated to generate noisy images characterized by their percentage phase coherence (Dakin et al., 2002). We used a total of four different phase coherence values (22%, 30%, 45%, and 55%), chosen based on behavioral pilot experiments, so overall behavioral performance spanned the psychophysical dynamic range. Specifically, the participants scored 52.1%, 64.7%, 85.2% and 98.7% in the mentioned coherence levels in the piloting. At each of the four phase coherence levels, we generated multiple frames for every image: the number of frames generated depended on the reaction time of the participants, as explained below. Different sets of participants were used for the actual and pilot experiments.

### EEG acquisition and Apparatus

We recorded EEG data from participants while they were performing the face categorization task. EEG data were acquired in an electrostatically shielded room using an ANT Neuro Amplifier (eego 64 EE-225) from 64 Ag/AgCl scalp electrodes and from three periocular electrodes placed below the left eye and at the left and right outer canthi. All channels were referenced to the left mastoid with input impedance <15k and chin ground. Data were sampled at 1000 Hz and a software-based 0.1-200 Hz bandpass filter was used to remove DC drifts, and high-frequency noise and 50 and 100 Hz (harmonic) notch filters were applied to minimize line noise. These filters were applied non-causally (using MATLAB filtfilt) to avoid phase-related distortions. We used Independent Component Analysis (ICA) to remove artefactual components in the signal. The components which were reflecting artefactual signals (eye movements, head movements) were removed based on ADJUST’s criteria (Mognon et al., 2011). Next, trials with strong eye movement or other movement artifacts were removed using visual inspection. On average, we kept 98.74%±1.5% artifact-free trials for any given condition.

We presented images on LCD monitor (BenQ XL2430, 24”, 144 Hz refresh rate, resolution of 1920 ×1080 pixels) and the stimulus presentation was controlled using custom-designed MATLAB codes and Psychtoolbox 3.0 (Brainard, 1997; Pelli, 1997). We presented stimuli at a distance of 60 cm to the participant, and each image subtended 8° × 8° of visual angle.

### Procedure

Participants performed a familiar vs. unfamiliar face categorization task by categorizing dynamically updating sequences of either familiar or unfamiliar face images in two recording sessions (Figure 1A). Image sequences were presented in rapid serial visual presentation (RSVP) fashion at a frame rate of 60 Hz frames per second (i.e. 16.67 ms per frame without gaps). Each trial consisted of a single sequence of up to 1.2 seconds (until response) with a series of images from the same stimulus (i.e., selection from either familiar or unfamiliar face categories) at one of the four possible phase coherence levels. Importantly, within each phase coherence level, the overall amount of noise remained unchanged, whereas the spatial distribution of the noise varied across individual frames such that different parts of the underlying image was revealed sequentially.

We instructed participants to fixate at the center of the monitor and respond as accurately and quickly as possible by pressing one of two keyboard keys (left and right arrow keys) to identify the image as familiar or unfamiliar using the right index and middle fingers, respectively. The mapping between familiar-unfamiliar categories and the two fingers were swapped from the first session to the next (counterbalanced across participants) and the data were collapsed across the two sessions before analyses. As soon as a response was given, the RSVP sequence stopped, followed by an inter-trial interval of 1–1.2 s (random with uniform distribution). The maximum time for the RSVP sequence was 1.2 secs. If participants failed to respond within the 1.2 secs period, the trial was marked as a no-choice trial and was excluded from further analysis. We had a total of 240 trials (i.e., 30 trials per perceptual category, familiar and unfamiliar, each at four phase coherence levels) during the experiment. Participants were naïve about the number and proportion of the face stimuli in categories. We presented six blocks of 36 trials each, and one block of 24 trials and participants had some resting time between the blocks. Each image from the image set was presented to the participants once in each session.

### Analysis

#### Decoding (MVPA) analysis

We decoded the information content of our conditions using Multivariate Pattern Analysis (MVPA) methods with Support Vector Machine (SVM) classifiers (Cortes et al., 1995). MVPA utilizes within-condition similarity of trials and their cross-condition dissimilarity to determine the information content of individual conditions. We trained an SVM classifier on the patterns of brain activity (from 64 EEG electrodes) from 90% of familiar (including personally familiar, famous, and self sub-categories) and 90% of unfamiliar trials, and then tested the trained classifier on the left-out 10% of trials from each category. The classification accuracy from categorization of the testing data shows whether there is information about familiarity in the neural signal. We only used the trials in which the participant *correctly* categorized the stimulus as familiar or unfamiliar. We repeated this procedure iteratively 10 times until all trials from the two categories were used in the testing of the classifier once (no trial was included both in the training and testing sets in a single run), hence 10-fold cross-validation, and averaged the classification accuracy across the 10 validation runs for each participant. To obtain the decoding accuracy through time, we down-sampled the EEG signals to 100 Hz and repeated the same classification procedure for every 10 ms time point from −100 to 600 ms relative to the onset of the stimulus, and from −500 to 100 ms relative to the response. This allowed us to assess the evolution of face familiarity information relative to the stimulus onset and response separately.

To investigate the potential differences in the temporal evolution of the sub-categories contained in the familiar category (i.e., famous, personally familiar and self), we additionally calculated the decoding accuracy for each sub-category separately. Note that the same decoding results obtained from decoding of familiar vs. unfamiliar categories were used here, only calculated separately for each sub-category of familiar faces. Finally, we averaged the decoding accuracies across participants and reported the group-level results.

We used random bootstrapping testing to evaluate the significance of the decoding accuracies at every time point for the group of participants. For every time point, this involved randomizing the labels of the familiar and unfamiliar trials 10,000 times and obtaining 10,000 decoding accuracies using the above procedure for each participant. Then we averaged the 10,000 decoding accuracies across (18) participants obtaining a single decoding accuracy for each of the 10,000 randomization for group-level analysis. For every time point, the p-value of the true group-averaged decoding accuracy was obtained as [1-p(10,000 randomly generated decoding accuracies which were surpassed by the corresponding true group-averaged decoding value)]. Since there is a different number of trials in each familiar sub-category, in the random bootstrapping, we maintained the same proportion of trials in each sub-category to preserve the original structure and generate an appropriate null distribution. We then corrected the *p* values for multiple comparisons across time (using MATLAB’s mafdr function at p<0.05). After the correction, the true decoding values with p < 0.05 were considered significantly above chance (e.g., 50%).

#### Brain-behavior correlation

To investigate if the decoding results could explain the observed behavioral face categorization results, we calculated the correlation between the decoding and the behavioral results using Spearman’s rank correlation. We calculated the correlation between a 16-element vector containing each participant’s behavioral accuracy for the four coherence levels of the four familiarity levels (i.e. Familiar, Famous, Self and Unfamiliar), and another vector with the same structure containing the decoding values from the same conditions (Karimi-Rouzbahani et al., 2020b). We repeated this procedure for every time point and each individual participant separately. Finally, we averaged the correlations across participants and reported the group-level results.

To determine the significance of the group-averaged correlations, the same bootstrapping procedure as described above was repeated at every time point by generating 10,000 random correlations after shuffling the elements of the 16-element behavioral vector. We repeated this procedure for every time point and each individual participant separately. Then we averaged the 10,000 random correlations across (18) participants obtaining a single correlation value for each of the 10,000 randomization for group-level analysis. For every time point, the p-value of the true group-averaged correlation was obtained as [1-p(10,000 randomly generated correlations which were surpassed by the corresponding true group-averaged correlation)]. We then corrected the *p* values for multiple comparisons across time (using MATLAB’s mafdr function at p<0.05). After the correction, the true correlation values with p < 0.05 were considered significantly above chance (i.e., 0).

#### Representational similarity analysis

Representational similarity analysis is used here for three purposes. First, to partial out the possible contributions of low-level image statistics to our decoding results, which is not directly possible in the decoding analysis (Supplementary Text). Second, to investigate possible coding strategies that the brain might have adopted which could explain our decoding, specifically, whether the brain was coding familiar versus unfamiliar faces, the different levels of familiarity or a combination of the superordinate and subordinate categories. Third, to measure the contribution of information from other brain areas to the representations of each given area (see Information flow analysis).

We constructed neural representational dissimilarity matrices (RDMs) by calculating the (*Spearman’s* rank) correlation between every possible representation obtained from every single presented image which resulted in a *correct* response (leading to a 240 by 240 RDM matrix if all images were categorized correctly, which was never the case for any participant). The matrices were constructed using signals from the electrodes over the whole brain as well as from peri-occipital and peri-frontal electrodes separately as explained later (Figures 4–6). We also constructed *image* RDMs for which we calculated the correlations between every possible pair of images which had generated the corresponding neural representations used in the neural RDMs (i.e. only from *correct* trials). Finally, to evaluate how much the neural RDMs coded the familiar vs. unfamiliar faces, familiar and unfamiliar faces separately, familiarity levels and each level of familiarity, we constructed different model RDMs. For examples, in the *Familiar-Unfamiliar* model RDM, the elements which corresponded to the correlations of familiar with familiar, or unfamiliar with unfamiliar, representations (and not their cross-correlations) were valued as 1, and the elements which corresponded to the cross-correlations between familiar and unfamiliar faces were valued as 0. The *Familiarity level* model, on the other hand, was filled with 0s (instead of 1s) for the representations which corresponded to the cross-correlations between different sub-categories of familiar faces (e.g. personally familiar vs. famous) with everything else being the same as the *Familiar-Unfamiliar* model RDM. Please note that the number of trials within all conditions of the RDM were down-sampled to the minimum number available for all conditions. This avoided potential difference across conditions as a result of unbalanced number of trials across conditions. To correlate the RDMs, we selected and reshaped the upper triangular elements of the RDMs (excluding the diagonal elements) into vector RDMs (or RDVs). To evaluate the correlation between the neural RDVs and the model RDVs, we used *Spearman’s* partial correlation in which we calculated the correlation between the neural and the model RDV while partialling out the image RDV as in equation (1):

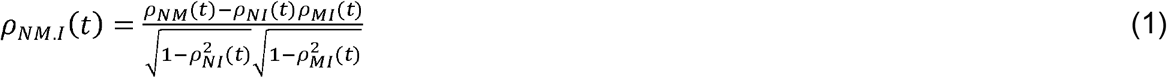

where *ρ* refers to Spearman correlation and *ρ_NM.I_* refers to the Spearman correlation between the neural and model RDVs after partialling out the image RDV. *N, M* and *I* respectively refer to Neural, Model and Image RDVs. As indicated in the equation, the partial correlation was calculated for every time point of the neural data (10 ms time steps), relative to the stimulus onset and response separately using the time-invariant model and image RDVs. To evaluate the significance of the partial correlations, we used a similar bootstrapping procedure as was used in decoding. However, here we randomized the elements of the model RDV 10,000 times (while keeping the number of ones and zeros equal to the original RDV) and calculated 10,000 random partial correlations. Finally, we compared the true partial correlation at every time point with the randomly generated partial correlations for the same time point and deemed it significant if it exceeded 95% of the random correlations (p < 0.05) after correcting for multiple comparisons.

#### Informational connectivity analysis

We developed a novel model-based method of information flow analysis to investigate how earlier information content of other brain areas contributes to the present-time information content of a given area. While several recent approaches have suggested for information flow analysis in the brain (Goddard et al., 2016; Karimi-Rouzbahani, 2018; Karimi-Rouzbahani et al., 2019), following the recent needs for these approaches in answering neuroscience questions (Anzellotti and Coutanche, 2018), none of the previously developed methods could answer the question of whether the transferred information was improving the representation of the target area in line with the behavioral task demands. Our proposed model, however, explicitly incorporates the specific aspects of behavioral goals or stimuli in its formulation and allows us to measure if the representations of target areas are shifted towards the behavioral/neural goals by the received information. An alternative would be that the incoming information from other areas are just epiphenomenal and are task-irrelevant. This new method can distinguish these alternatives.

Accordingly, we split the EEG electrodes in two groups, each with 16 electrodes: peri-frontal and peri-occipital (Figure 4A) to see how familiarity information is (if at all) transferred between these areas that can be broadly categorized as “cognitive” and “sensory” brain areas, respectively. We calculated the neural RDMs for each area separately and calculated the correlation between the neural RDV and the model RDV, partialling out the image RDM from the correlation (as explained in equation (1)). This resulted in a curve when calculating the partial correlation at every time point in 10 ms intervals (see the solid lines in Figure 4B). Note that the partial correlation curve for the peri-frontal area could have received contributions from the present and earlier representations of the same area (i.e., the latter being imposed by our sequential stimulus presentation). It could also have received contributions from earlier peri-occipital representations through information flow from peri-occipital to the peri-frontal area. To measure this potential contribution, we partialled out the earlier peri-occipital representations in calculation of the partial correlation between peri-frontal and model RDVs and calculated the difference between the former and the latter partial correlations as feed-forward information flow according to equation (2):

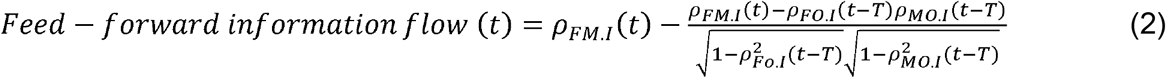

where *ρ_FM.I_* refers to the partial correlation between the peri-frontal and the model RDV, *ρ_FO.I_* the partial correlation between peri-frontal and peri-occipital RDVs and *ρ_MO.I_* the partial correlation between the peri-occipital and model RDVs. Please note that the image RDV is partialled out from all pairwise correlations to remove its effect in the analysis, so the subscript *I* and the term “partial”. This determines the contribution of earlier peri-occipital representations to the present peri-frontal areas which we called “feed-forward information flow” (as indicated by the brown shades in Figure 4). To determine the contribution of the peri-frontal representations in modulating the peri-occipital representations, we used equation (3):

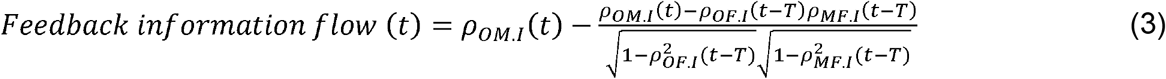

with the same notations as in equation (2). Accordingly, equation (3) determines the contribution of earlier peri-frontal representations in directing the peri-occipital representations towards the model RDV, namely ‘feedback information flow’. In equations (2) and (3), the delay time (T) was 30ms, which was selected based on previously reported delay times between the peri-occipital and peri-frontal areas in visual processing (Foxe and Simpson, 2002). To that end, five earlier RDVs were averaged (5 time points centered on −30ms) leading to an average delay time of 30ms.

Finally, to characterize the information flow dynamics between the peri-occipital and peri-frontal areas, we calculated the difference between the feed-forward and feedback contribution of information flows. This allowed us to investigate the transaction of targeted information between the brain areas aligned to the stimulus onset and response. We repeated the same procedure using the Familiar-Unfamiliar as well as Familiarity level models to see if they differed. We validated the proposed informational connectivity method using simulated well-controlled dataset (Supplementary Figure 6). We determined the significance of the partial correlations using the above-explained random bootstrapping procedure. We determined the significance of the differences between partial correlations (the shaded areas in Figure 4 and the lines in panel C) and the differences in the feed-forward and feedback contribution of information using Wilcoxon’s signed-rank test using p < 0.05 threshold for significance after correction for multiple comparisons (using Matlab mafdr).

## Acknowledgements

We would like to thank Ali Yoonessi for supporting our electroencephalography data collection in his lab. We would also like to appreciate Chris I Baker, Mark Williams, Erin Goddard and Hamed Nili for providing their valuable feedback on the manuscript. This research was funded by UK Royal Society’s Newton International Fellowship NIF\R1\192608 to H.K.R. and MRC intramural funding SUAG/052/G101400 to A.W.

## Supplementary Materials

**Supplementary Figure 1.**
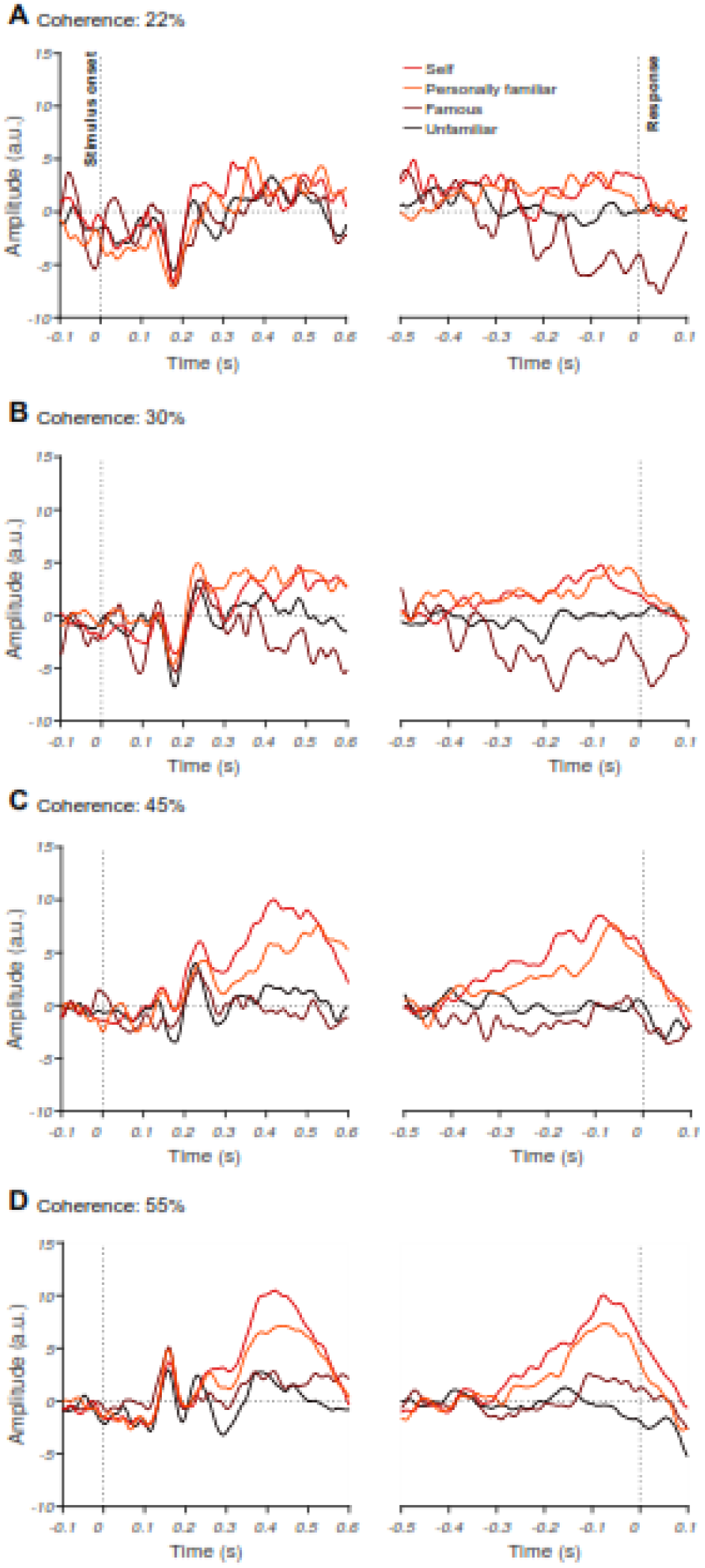
The effect of familiarity and sensory evidence on event-related potentials (ERPs). Averaged ERPs for 22% (A), 30% (B), 45% (C) and 55% (D) phase coherence levels and four face categories across all participants for an electrode at a centroparietal site (CP2). Note that the left panels show stimulus-aligned ERPs while the right panel shows response-aligned ERPs. The differences between levels of familiarity were more pronounced at later stages of stimulus processing around 400 ms post-stimulus onset and <100 ms before the response was given by the participant in the stimulus- and response-aligned analyses, respectively.

**Supplementary Figure 2.**
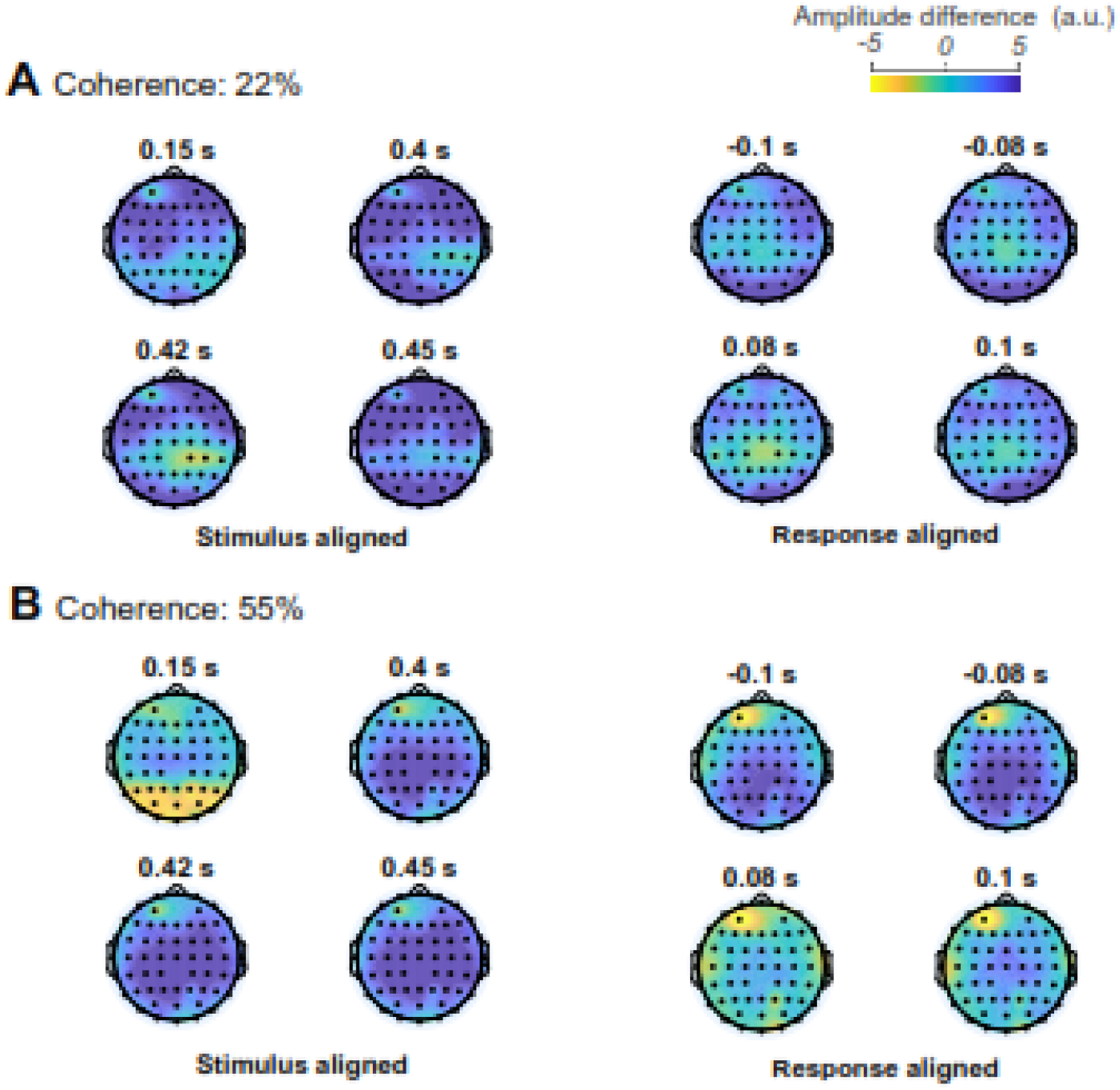
Familiarity information on the whole-brain event-related potentials (ERPs). The topographic maps show the difference in ERPs between unfamiliar and the average of the three familiar face categories (i.e. unfamiliar-average of unfamiliar categories) at specific time points averaged across participants. The time points were chosen based on the results from Figure 2, when the ERPs were significantly (p<0.05) higher in the 55% vs. 22% coherence levels. Note that the left panels show stimulus-aligned ERPs while the right panel shows response-aligned ERPs.

**Supplementary Figure 3.**
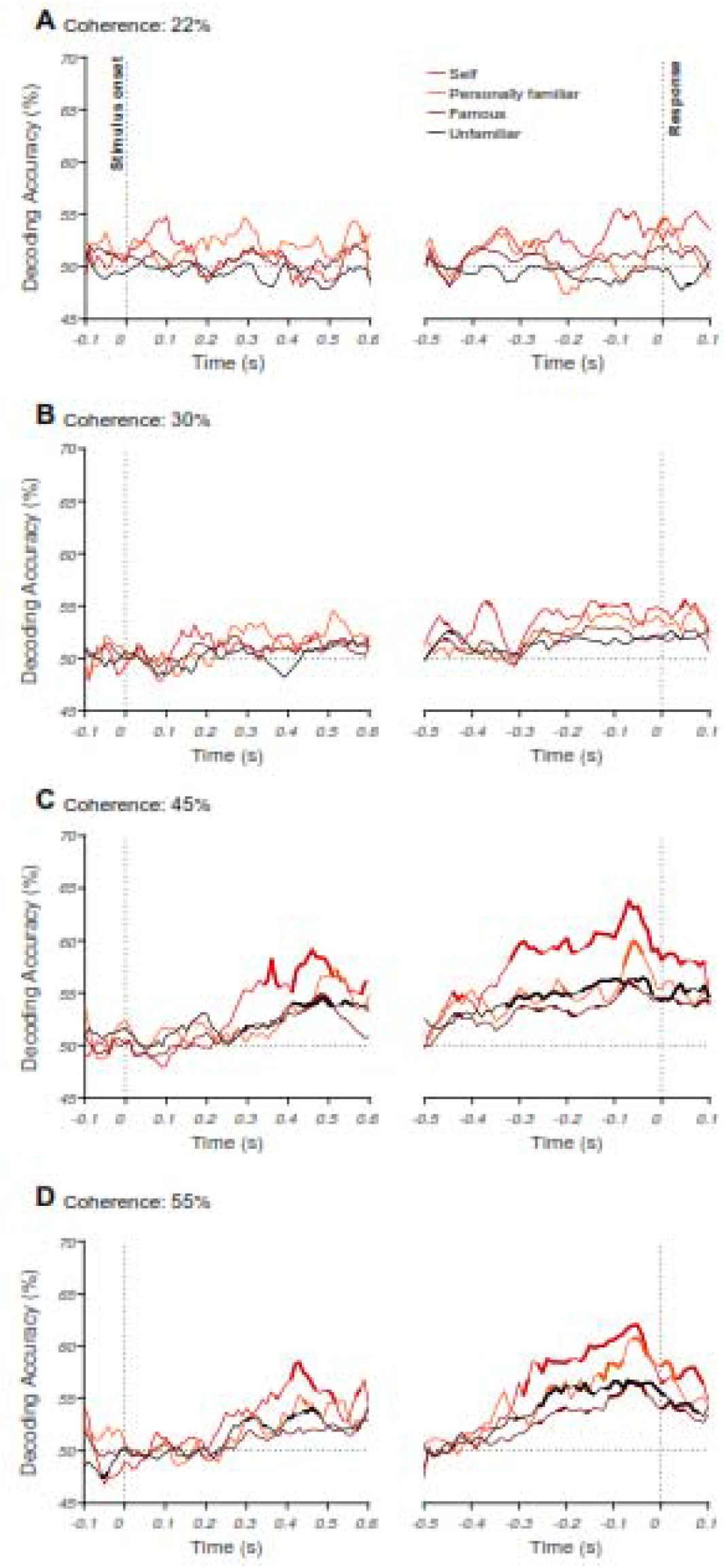
Decoding of face familiarity from EEG signals. Time course of decoding accuracy for familiar versus unfamiliar faces from EEG signals for four different phase coherence levels (22% (A), 30% (B), 45% (C), and 55% (D)). The chance accuracy is 50%. Thickened lines indicate the time points when the accuracy was significantly above chance level (sign rank test, FDR corrected across time, p<0.05). The left panels show the results for stimulus-aligned analysis while the right panels show the results for response-aligned analysis (averaged over 18 participants).

### Supplementary Text: Low-level image statistics do not explain the separation of familiar from unfamiliar faces

Although, we did equalize the frequency content, pixel intensities and contrast of the images of our dataset (see *methods*), but we checked whether there are other low-level differences by creating a model representational dissimilarity matrix (RDM) for each of the categories under different phrase coherences. Briefly, *neural* RDMs are constructed by calculating the correlations (or dissimilarities) of the *brain* response to different face stimuli to give an abstract representation of information encoding in the brain. We also construct a *low-level feature* RDM, for which we calculate the correlations between images corresponding to each brain response. *Model* RDMs *predicted* representations in the brain (see *Methods*). The model RDMs were created for discriminating (1) familiar from unfamiliar (Supplementary Figure 4A) and also (2) the familiarity levels from one another (Supplementary Figure 4B). We then computed partial Spearman’s correlations between one of the models and neural RDMs for every time point and participant, while partialling out (Supplementary Figure 4)/not partialling out (Supplementary Figure 5) low-level feature model RDM.

This analysis revealed the emergence of familiarity representation (familiar vs. unfamiliar faces) at around 270 ms post-stimulus for the highest coherence level (55%, Supplementary Figure 4A). The onset of significant representation is slightly later for lower coherence levels (e.g., 45%, Supplementary Figure 4A), which may suggest the need for additional processing time required to evaluate the sensory evidence. Interestingly, while the dynamics of familiarity level representations also showed gradual accumulation of information (Supplementary Figure 4B), especially for the 45% and 55% coherence, the correlation values are generally higher for the model of familiarity level compared to familiar-unfamiliar (c.f. Supplementary Figure 4A). This suggests that there might be well-established neural mechanisms in the brain that discriminate levels of familiarity so strongly that is not suppressed/dominated by the task (i.e. here familiar-unfamiliar) or the response of the participants. This could also be supported by the observation that, as opposed to the familiar-unfamiliar representations, for which the 55% coherence provided the most information (at least in the stimulus-aligned analysis), the familiarity level representations provided their highest information in lower coherence levels such as 45% (in both stimulus- and response-aligned analyses) and 30% or even 22% in the response-aligned analysis. Note that participants’ task and response could have also potentially contributed to the analysis of face familiarity model as those factors matched the familiar-unfamiliar model used in Supplementary Figure 4A.

**Supplementary Figure 4.**
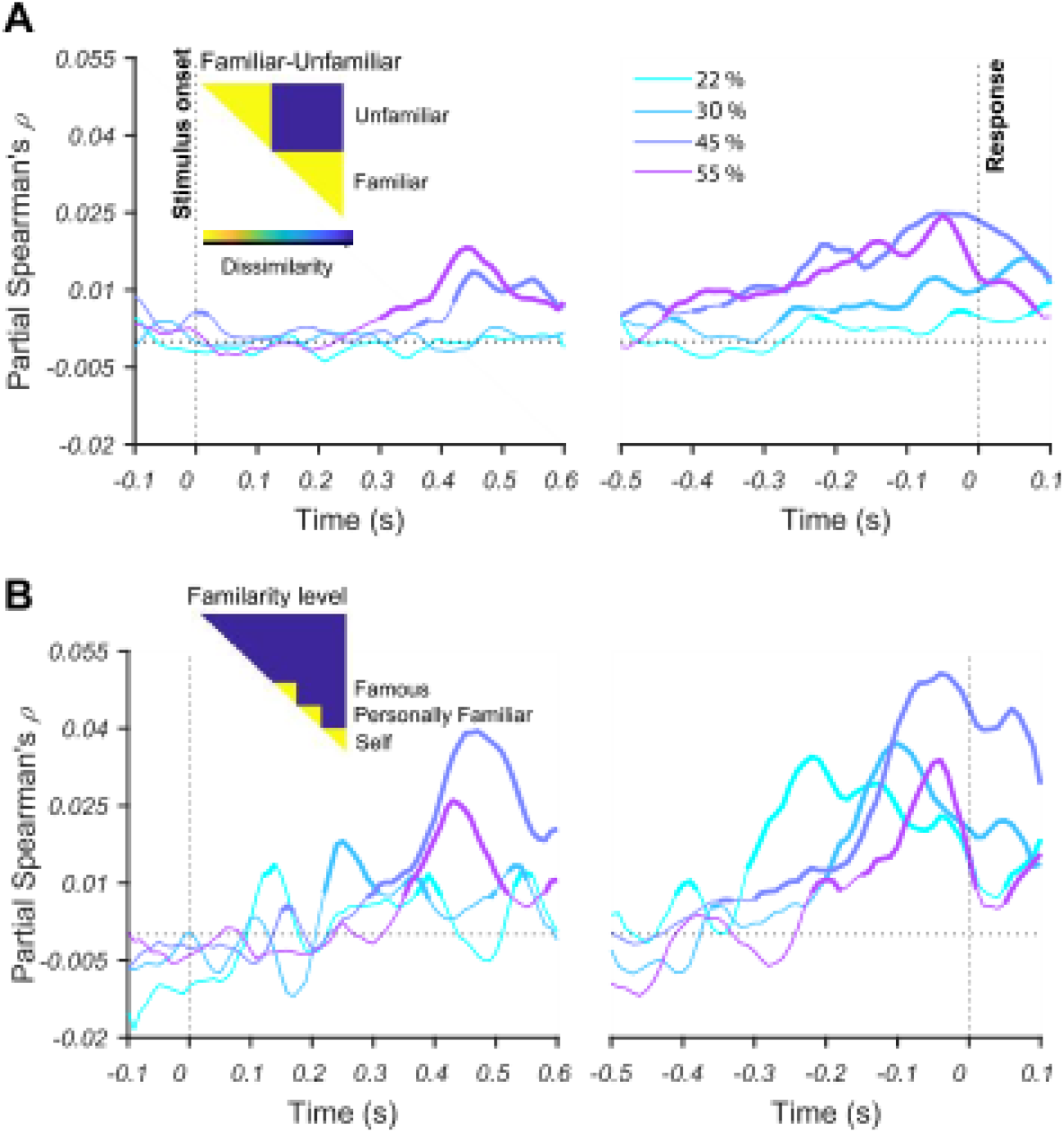
Representations of face familiarity and categories revealed by RSA. Time course of Spearman’s correlations between neural RDMs and model RDM (shown as insets) for (A) face familiarity; and (B) face familiarity levels, famous, self and personally familiar faces, after partialling out contributions from low-level features (see Methods). Each colored trace shows the correlations over time for one phase coherence level. Thickened lines indicate time points where the correlation is significant (sign permutation test, FDR-corrected significance level at p□<□0.05), and black horizontal dotted lines indicate 0 correlation. The left panels show the results for stimulus-aligned analysis while the right panels represent the results for response-aligned analysis.

Apart from a small difference in absolute decoding rates, the dynamics of neural representations were similar when not partialling out the low-level feature model RDM (Supplementary Figure 5), presenting the ramping up of information, with earlier and most mounting trends for highest coherence levels (i.e. 45% and 55%). The similar patterns of neural information decoding between the correlation patterns with and without the low-level feature model suggest that low-level image statistics may only play a minor role in driving the observed decoding analyses. Nonetheless, we partialled out the low-level feature model in all the following RSA-based analyses to avoid their potential contribution to the results.

**Supplementary Figure 5.**
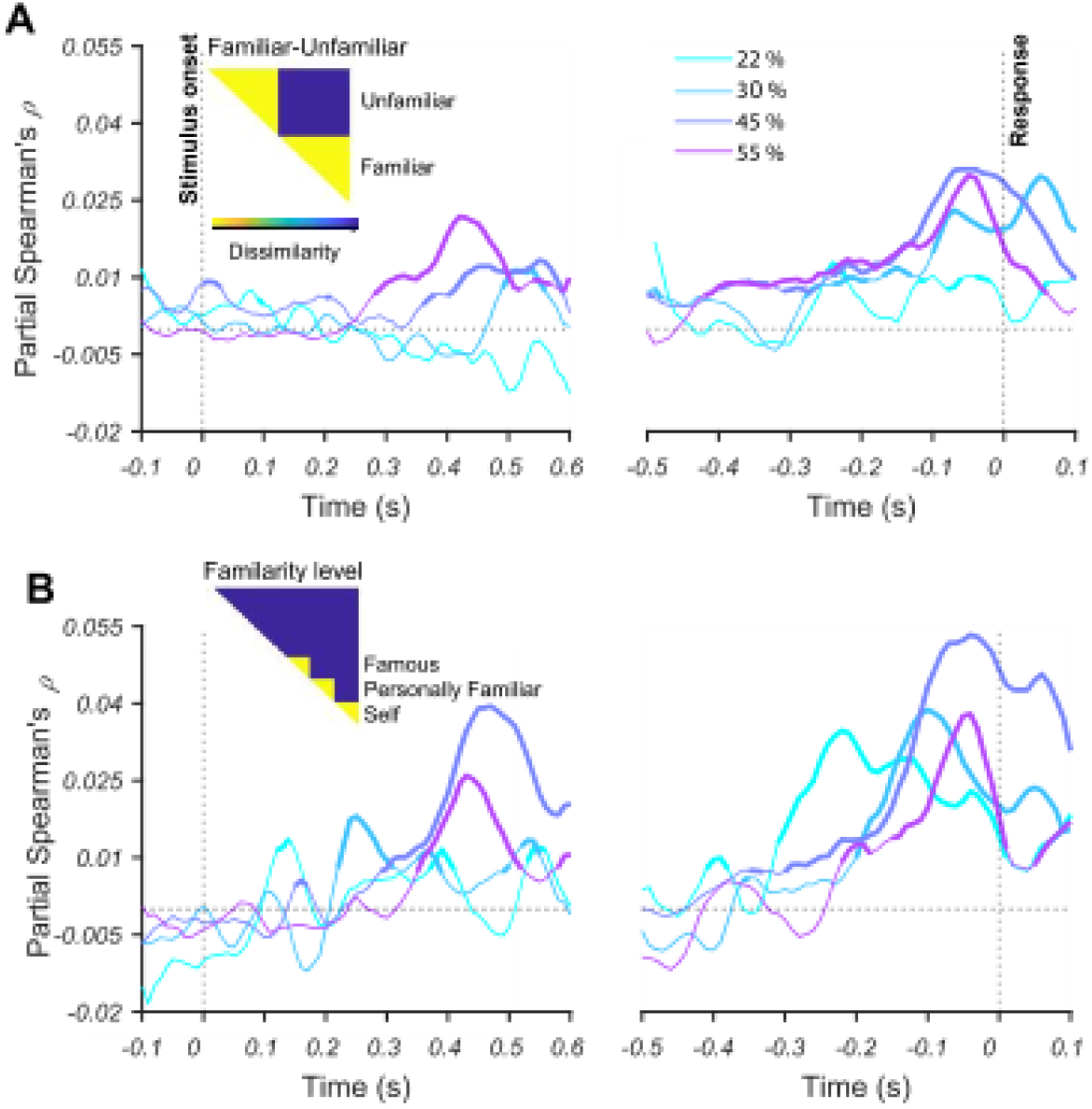
Representations of face familiarity and categories revealed by RSA. Time course of Spearman’s correlations between neural RDMs and model RDM (shown as insets) for (A) face familiarity; and (B) face familiarity levels, famous, self and personally familiar faces, before partialling out contributions from low-level features (see Methods). Each colored trace shows the correlations over time for one phase coherence level. Thickened lines indicate time points where the correlation is significant (sign permutation test, FDR-corrected significance level at p□<□0.05), and black horizontal dotted lines indicate 0 correlation. The left panels show the results for stimulus-aligned analysis while the right panels represent the results for response-aligned analysis. Note that the correlation values are higher compared to the results after partialling out contributions from low-level features (see Supplementary Figure 4).

**Supplementary Figure 6.**
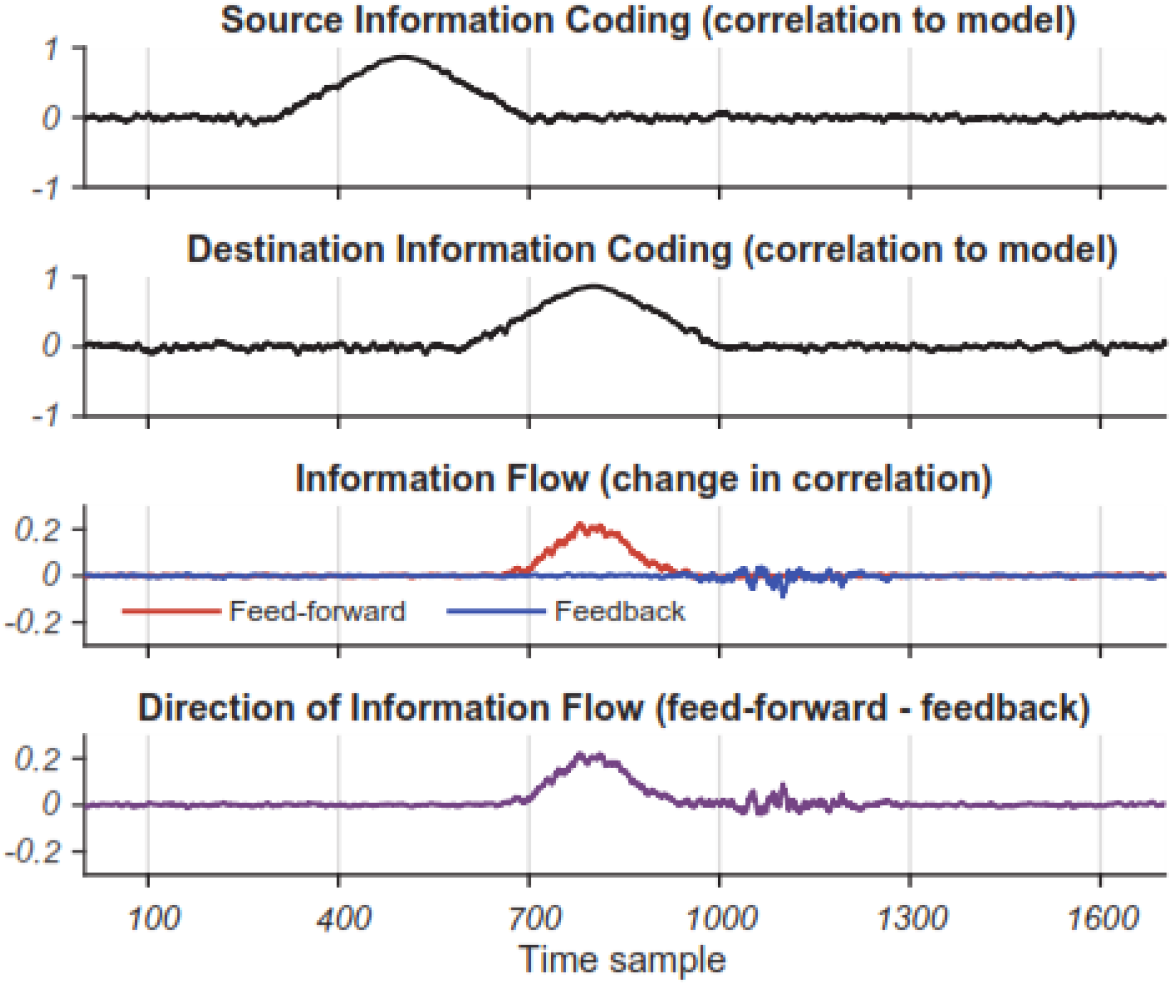
Simulation of informational connectivity and its measurement using the proposed connectivity analysis. Feed-forward and feedback refer to information flow from source to destination and vice versa, respectively. For this stimulation, we initially generated three static 16*16 RDMs, one for source, one for destination and one for model, all similar to the familiar-unfamiliar RDMs we presented throughout the manuscript (e.g. Figure 4A). Next, by adding varying levels of uniform noise, in the range between 0 and 1, to the artificially generated source and destination static RDMs, we generated temporally changing dynamics of information coding (i.e. measured as how much they correlated with the desired model RDM) in those RDMs across time samples. The peak of information coding in the destination RDM was designed to appear ***300*** samples after the coding in the source RDM, so that it simulates flow of information in the feed-forward direction. Finally, we applied our connectivity analysis to these data to check if it could detect the information flow from source to destination area. The two top panels show the correlation between temporally varying simulated source and destination RDMs and the model RDM. Third panel from top shows the amount of feed-forward and feedback information and the bottom panel shows their difference as measured by our informational connectivity analysis. Results show a clear feed-forward (from source to destination) information flow and almost zero information flow in the feedback (from destination to source) direction, which peaks almost simultaneously with the information peak in the destination area. This result suggests that our informational connectivity detects the simulated connectivity in the correct direction and temporal dynamics.

## Footnotes

1 Here we use the terms “peri-occipital” and “peri-frontal” to refer broadly to groups of electrodes selected from posterior and anterior parts of the EEG cap, respectively (as indicated in Figure 5).

2 http://mmlab.ie.cuhk.edu.hk/proiects/CelebA.html https://megapixels.ee/datasets/msceleb/

3 https://github.com/Masoud-Ghodrati/face_familiarity

## References

Ambrus, Géza Gergely, Daniel Kaiser, Radoslaw Martin Cichy, and Gyula Kovács. 2019. “The Neural Dynamics of Familiar Face Recognition.” Cerebral Cortex 29 (11): 4775–4784.

Anzellotti, Stefano, and Marc N. Coutanche. 2018. “Beyond Functional Connectivity: Investigating Networks of Multivariate Representations.” Trends in Cognitive Sciences 22 (3): 258–269.

Bar, Moshe, Karim S. Kassam, Avniel Singh Ghuman, Jasmine Boshyan, Annette M. Schmid, Anders M. Dale, Matti S. Hämäläinen, Ksenija Marinkovic, Daniel L. Schacter, and Bruce R. Rosen. 2006. “Top-down Facilitation of Visual Recognition.” Proceedings of the National Academy of Sciences 103 (2): 449–454.

Basti, Alessio, Hamed Nili, Olaf Hauk, Laura Marzetti, and Richard Henson. 2020. “Multi-dimensional Connectivity: A Conceptual and Mathematical Review.” Neuroimage 117179.

Besson, Gabriel, Gladys Barragan-Jason, Simon J. Thorpe, Michèle Fabre-Thorpe, Sébastien Puma, Mathieu Ceccaldi, and Emmanuel J. Barbeau. 2017. “From Face Processing to Face Recognition: Comparing Three Different Processing Levels.” Cognition 158: 33–43.

Brainard, David H. 1997. “The Psychophysics Toolbox.” Spatial Vision 10 (4): 433–436.

Brown, M. W., and P. J. Banks. 2015. “In search of a recognition memory engram.” Neuroscience & Biobehavioral Reviews 50: 12–28.

Caharel, Stephanie, Stephane Poiroux, Christian Bernard, Florence Thibaut, Robert Lalonde, and Mohamed Rebai. 2002. “ERPs associated with familiarity and degree of familiarity during face recognition.” International Journal of Neuroscience 112(12): 1499–1512.

Chen, Yao, Susana Martinez-Conde, Stephen L. Macknik, Yulia Bereshpolova, Harvey A. Swadlow, and Jose-Manuel Alonso. 2008. “Task Difficulty Modulates the Activity of Specific Neuronal Populations in Primary Visual Cortex.” Nature Neuroscience 11 (8): 974.

Clarke, Alex, Barry J. Devereux, and Lorraine K. Tyler. 2018. “Oscillatory Dynamics of Perceptual to Conceptual Transformations in the Ventral Visual Pathway.” Journal of Cognitive Neuroscience 30 (11): 1590–1605.

Collins, Elliot, Amanda K. Robinson, and Marlene Behrmann. 2018. “Distinct Neural Processes for the Perception of Familiar versus Unfamiliar Faces along the Visual Hierarchy Revealed by EEG.” NeuroImage 181: 120–131.

Collins, Jessica A., and Ingrid R. Olson. 2014. “Beyond the FFA: the role of the ventral anterior temporal lobes in face processing.” Neuropsychologia 61: 65–79.

Dakin, S. C., R. F. Hess, T. Ledgeway, and R. L. Achtman. 2002. “What Causes Non-Monotonic Tuning of FMRI Response to Noisy Images?” Current Biology 12 (14): R476–R477.

Davies-Thompson, Jodie, and Timothy J. Andrews. 2012. “Intra-and interhemispheric connectivity between face-selective regions in the human brain.” Journal of Neurophysiology 108 (11): 3087–3095.

Delorme, Arnaud, Guillaume A. Rousselet, Marc J.-M. Macé, and Michele Fabre-Thorpe. 2004. “Interaction of Top-down and Bottom-up Processing in the Fast Visual Analysis of Natural Scenes.” Cognitive Brain Research 19 (2): 103–113.

Dobs, Katharina, Leyla Isik, Dimitrios Pantazis, and Nancy Kanwisher. 2019. “How Face Perception Unfolds over Time.” Nature Communications 10 (1): 1–10.

Duchaine, Brad, and Galit Yovel. 2015. “A revised neural framework for face processing.” Annual review of vision science 1: 393–416.

Ellis, Hadyn D., John W. Shepherd, and Graham M. Davies. 1979. “Identification of Familiar and Unfamiliar Faces from Internal and External Features: Some Implications for Theories of Face Recognition.” Perception 8 (4): 431–439.

Ethofer, Thomas, Markus Gschwind, and Patrik Vuilleumier. 2011. “Processing social aspects of human gaze: a combined fMRI-DTI study.” Neuroimage 55 (1): 411–419.

Fan, Xiaoxu, Fan Wang, Hanyu Shao, Peng Zhang, and Sheng He. 2020. “The Bottom-up and Top-down Processing of Faces in the Human Occipitotemporal Cortex.” ELife 9: e48764.

Felleman, Daniel J., and DC Essen Van. 1991. “Distributed Hierarchical Processing in the Primate Cerebral Cortex.” Cerebral Cortex (New York, NY: 1991) 1 (1): 1–47.

Fenske, Mark J., Elissa Aminoff, Nurit Gronau, and Moshe Bar. 2006. “Top-down Facilitation of Visual Object Recognition: Object-Based and Context-Based Contributions.” Progress in Brain Research 155: 3–21.

Foxe, John J., and Gregory V. Simpson. 2002. “Flow of Activation from V1 to Frontal Cortex in Humans.” Experimental Brain Research 142 (1): 139–150.

Gilbert, Charles D., and Mariano Sigman. 2007. “Brain States: Top-down Influences in Sensory Processing.” Neuron 54 (5): 677–696.

Gilbert, Charles D., and Wu Li. 2013. “Top-down Influences on Visual Processing.” Nature Reviews Neuroscience 14 (5): 350–363.

Gobbini, M. Ida, Ellen Leibenluft, Neil Santiago, and James V. Haxby. 2004. “Social and Emotional Attachment in the Neural Representation of Faces.” Neuroimage 22 (4): 1628–1635.

Goddard, Erin, Thomas A. Carlson, and Alexandra Woolgar. 2019. “Spatial and Feature-Selective Attention Have Distinct Effects on Population-Level Tuning.” BioRxiv, 530352.

Goddard, Erin, Thomas A. Carlson, Nadene Dermody, and Alexandra Woolgar. 2016. “Representational Dynamics of Object Recognition: Feedforward and Feedback Information Flows.” Neuroimage 128: 385–397.

Gregoriou, Georgia G., Stephen J. Gotts, Huihui Zhou, and Robert Desimone. 2009. “High-Frequency, Long-Range Coupling between Prefrontal and Visual Cortex during Attention.” Science 324 (5931): 1207–1210.

Hanks, Timothy D., and Christopher Summerfield. 2017. “Perceptual Decision Making in Rodents, Monkeys, and Humans.” Neuron 93 (1): 15–31.

Hebart, Martin N., Brett B. Bankson, Assaf Harel, Chris I. Baker, and Radoslaw M. Cichy. 2018. “The representational dynamics of task and object processing in humans.” Elife 7: e32816.

Henson, Richard N., Elias Mouchlianitis, William J. Matthews, and Sid Kouider. 2008. “Electrophysiological Correlates of Masked Face Priming.” Neuroimage 40 (2): 884–895.

Huang, Wanyi, Xia Wu, Liping Hu, Lei Wang, Yulong Ding, and Zhe Qu. 2017. “Revisiting the Earliest Electrophysiological Correlate of Familiar Face Recognition.” International Journal of Psychophysiology 120: 42–53.

Hupé, J. M., A. C. James, B. R. Payne, S. G. Lomber, P. Girard, and J. Bullier. 1998. “Cortical Feedback Improves Discrimination between Figure and Background by V1, V2 and V3 Neurons.” Nature 394 (6695): 784–787.

Johnson, Matthew R., Karen J. Mitchell, Carol L. Raye, Mark D’Esposito, and Marcia K. Johnson. 2007. “A Brief Thought Can Modulate Activity in Extrastriate Visual Areas: Top-down Effects of Refreshing Just-Seen Visual Stimuli.” Neuroimage 37 (1): 290–299.

Karimi-Rouzbahani, Hamid, Alexandra Woolgar, and Anina N. Rich. 2020a. “Neural signatures of vigilance decrements predict behavioral errors before they occur.” bioRxiv.

Karimi-Rouzbahani, Hamid, Ehsan Vahab, Reza Ebrahimpour, and Mohammad Bagher Menhaj. 2019. “Spatiotemporal Analysis of Category and Target-Related Information Processing in the Brain during Object Detection.” Behavioral Brain Research 362: 224–239.

Karimi-Rouzbahani, Hamid, Mozhgan Shahmohammadi, Ehsan Vahab, Saeed Setayeshi, and Thomas Carlson. 2020b. “Temporal codes provide additional category-related information in object category decoding: a systematic comparison of informative EEG features.” bioRxiv.

Karimi-Rouzbahani, Hamid, Nasour Bagheri, and Reza Ebrahimpour. 2017a. “Average activity, but not variability, is the dominant factor in the representation of object categories in the brain.” Neuroscience 346: 14–28.

Karimi-Rouzbahani, Hamid, Nasour Bagheri, and Reza Ebrahimpour. 2017b. “Hard-Wired Feed-Forward Visual Mechanisms of the Brain Compensate for Affine Variations in Object Recognition.” Neuroscience 349: 48–63.

Karimi-Rouzbahani, Hamid, Nasour Bagheri, and Reza Ebrahimpour. 2017c. “Invariant Object Recognition Is a Personalized Selection of Invariant Features in Humans, Not Simply Explained by Hierarchical Feed-Forward Vision Models.” Scientific Reports 7 (1): 1–24.

Karimi-Rouzbahani, Hamid. 2018. “Three-Stage Processing of Category and Variation Information by Entangled Interactive Mechanisms of Peri-Occipital and Peri-Frontal Cortices.” Scientific Reports 8 (1): 1–22.

Kaufmann, Jürgen M., Stefan R. Schweinberger, and A. Mike Burton. 2009. “N250 ERP Correlates of the Acquisition of Face Representations across Different Images.” Journal of Cognitive Neuroscience 21 (4): 625–641.

Kay, Kendrick N., and Jason D. Yeatman. 2017. “Bottom-up and Top-down Computations in Word-and Face-Selective Cortex.” Elife 6: e22341.

Kelly, Simon P., and Redmond G. O’Connell. 2013. “Internal and External Influences on the Rate of Sensory Evidence Accumulation in the Human Brain.” Journal of Neuroscience 33 (50): 19434–19441.

Kietzmann, Tim C., Courtney J. Spoerer, Lynn Sörensen, Radoslaw M. Cichy, Olaf Hauk, and Nikolaus Kriegeskorte. 2019. “Recurrence Required to Capture the Dynamic Computations of the Human Ventral Visual Stream.” ArXiv Preprint ArXiv:1903.05946.

Kovács, Gyula. 2020 “Getting to Know Someone: Familiarity, Person Recognition, and Identification in the Human Brain.” Journal of Cognitive Neuroscience 1–19.

Kramer, Robin SS, Andrew W. Young, and A. Mike Burton. 2018. “Understanding Face Familiarity.” Cognition 172: 46–58.

Lamme, Victor AF, and Pieter R. Roelfsema. 2000. “The Distinct Modes of Vision Offered by Feedforward and Recurrent Processing.” Trends in Neurosciences 23 (11): 571–579.

Lamme, Victor AF, Karl Zipser, and Henk Spekreijse. 2002. “Masking Interrupts Figure-Ground Signals in V1.” Journal of Cognitive Neuroscience 14 (7): 1044–1053.

Landi, Sofia M., and Winrich A. Freiwald. 2017. “Two Areas for Familiar Face Recognition in the Primate Brain.” Science 357 (6351): 591–595.

Lee, Tai Sing, and David Mumford. 2003. “Hierarchical Bayesian Inference in the Visual Cortex.” JOSA A 20 (7): 1434–1448.

Leibenluft, Ellen, M. Ida Gobbini, Tara Harrison, and James V. Haxby. 2004. “Mothers’ Neural Activation in Response to Pictures of Their Children and Other Children.” Biological Psychiatry 56 (4): 225–232.

Leppänen, Jukka M., and Charles A. Nelson. 2009. “Tuning the developing brain to social signals of emotions.” Nature Reviews Neuroscience 10 (1): 37–47.

Mechelli, Andrea, Cathy J. Price, Karl J. Friston, and Alumit Ishai. 2004. “Where Bottom-up Meets Top-down: Neuronal Interactions during Perception and Imagery.” Cerebral Cortex 14 (11): 1256–1265.

Mognon, Andrea, Jorge Jovicich, Lorenzo Bruzzone, and Marco Buiatti. 2011. “ADJUST: An Automatic EEG Artifact Detector Based on the Joint Use of Spatial and Temporal Features.” Psychophysiology 48 (2): 229–240.

Mohsenzadeh, Yalda, Sheng Qin, Radoslaw M. Cichy, and Dimitrios Pantazis. 2018. “Ultra-Rapid Serial Visual Presentation Reveals Dynamics of Feedforward and Feedback Processes in the Ventral Visual Pathway.” Elife 7: e36329.

Norman, Kenneth A., Sean M. Polyn, Greg J. Detre, and James V. Haxby. 2006. “Beyond mind-reading: multi-voxel pattern analysis of fMRI data.” Trends in cognitive sciences 10 (9): 424–430.

Pelli, Denis G. 1997. “The VideoToolbox Software for Visual Psychophysics: Transforming Numbers into Movies.” Spatial Vision 10 (4): 437–442.

Philiastides, Marios G., and Paul Sajda. 2006. “Temporal Characterization of the Neural Correlates of Perceptual Decision Making in the Human Brain.” Cerebral Cortex 16 (4): 509–518.

Philiastides, Marios G., Roger Ratcliff, and Paul Sajda. 2006. “Neural Representation of Task Difficulty and Decision Making during Perceptual Categorization: A Timing Diagram.” Journal of Neuroscience 26 (35): 8965–8975.

Praß, Maren, Cathleen Grimsen, Martina König, and Manfred Fahle. 2013. “Ultra Rapid Object Categorization: Effects of Level, Animacy and Context.” PLoS One 8 (6).

Pratte, Michael S., Sam Ling, Jascha D. Swisher, and Frank Tong. 2013. “How Attention Extracts Objects from Noise.” Journal of Neurophysiology 110 (6): 1346–1356.

Ramon, Meike, and Maria Ida Gobbini. 2018. “Familiarity Matters: A Review on Prioritized Processing of Personally Familiar Faces.” Visual Cognition 26 (3): 179–195.

Ramon, Meike, Luca Vizioli, Joan Liu-Shuang, and Bruno Rossion. 2015. “Neural Microgenesis of Personally Familiar Face Recognition.” Proceedings of the National Academy of Sciences 112 (35): E4835–E4844.

Ress, David, Benjamin T. Backus, and David J. Heeger. 2000. “Activity in Primary Visual Cortex Predicts Performance in a Visual Detection Task.” Nature Neuroscience 3 (9): 940–945.

Schweinberger, Stefan R., Esther C. Pickering, Ines Jentzsch, A. Mike Burton, and Jürgen M. Kaufmann. 2002. “Event-Related Brain Potential Evidence for a Response of Inferior Temporal Cortex to Familiar Face Repetitions.” Cognitive Brain Research 14 (3): 398–409.

Shadlen, Michael N., and William T. Newsome. 2001. “Neural Basis of a Perceptual Decision in the Parietal Cortex (Area LIP) of the Rhesus Monkey.” Journal of Neurophysiology 86 (4): 1916–1936.

Spacek, Martin A., Gregory Born, Davide Crombie, Steffen A. Katzner, and Laura Busse. 2019. “Robust Effects of Cortical Feedback on Thalamic Firing Mode during Naturalistic Stimulation.” BioRxiv, 776237.

Sugiura, Motoaki, Carlos Makoto Miyauchi, Yuka Kotozaki, Yoritaka Akimoto, Takayuki Nozawa, Yukihito Yomogida, Sugiko Hanawa, Yuki Yamamoto, Atsushi Sakuma, and Seishu Nakagawa. 2015. “Neural Mechanism for Mirrored Self-Face Recognition.” Cerebral Cortex 25 (9): 2806–2814.

Summerfield, Christopher, and Tobias Egner. 2009. “Expectation (and attention) in visual cognition.” Trends in cognitive sciences 13 (9): 403–409.

Summerfield, Christopher, Tobias Egner, Matthew Greene, Etienne Koechlin, Jennifer Mangels, and Joy Hirsch. 2006. “Predictive codes for forthcoming perception in the frontal cortex.” Science 314 (5803): 1311–1314.

Supèr, Hans, Henk Spekreijse, and Victor AF Lamme. 2001. “Two Distinct Modes of Sensory Processing Observed in Monkey Primary Visual Cortex (V1).” Nature Neuroscience 4 (3): 304–310.

Taylor, Margot J., Marie Arsalidou, Sarah J. Bayless, Drew Morris, Jennifer W. Evans, and Emmanuel J. Barbeau. 2009. “Neural Correlates of Personally Familiar Faces: Parents, Partner and Own Faces.” Human Brain Mapping 30 (7): 2008–2020.

Visconti, M. di Oleggio Castello, and M. I. Gobbini. 2015. “Familiar Face Detection in 180 Ms.” PloS One 10 (8): e0136548–e0136548.

Wiese, Holger, Simone C. Tüttenberg, Brandon T. Ingram, Chelsea YX Chan, Zehra Gurbuz, A. Mike Burton, and Andrew W. Young. 2019. “A robust neural index of high face familiarity.” Psychological science 30 (2): 261–272.

Woolgar, Alexandra, Adam Hampshire, Russell Thompson, and John Duncan. 2011. “Adaptive Coding of Task-Relevant Information in Human Frontoparietal Cortex.” Journal of Neuroscience 31 (41): 14592–14599.

Woolgar, Alexandra, Soheil Afshar, Mark A. Williams, and Anina N. Rich. 2015. “Flexible Coding of Task Rules in Frontoparietal Cortex: An Adaptive System for Flexible Cognitive Control.” Journal of Cognitive Neuroscience 27 (10): 1895–1911.

Young, Andrew W., and A. Mike Burton. 2018. “Are We Face Experts?” Trends in Cognitive Sciences 22 (2): 100–110.

